# Paradoxical ventral tegmental area GABA signaling drives enhanced morphine reward after adolescent nicotine

**DOI:** 10.1101/2025.03.09.642288

**Authors:** Ruthie E. Wittenberg, Sanghee Yun, Kechun Yang, Olivia K. Swanson, Shannon L. Wolfman, Lorianna M. Colón, Amelia J. Eisch, John A. Dani

## Abstract

**Background:** An important yet poorly understood risk factor for opioid use disorder is adolescent nicotine use. We investigated the neural mechanisms underlying this understudied interaction.

**Methods:** Male and female adolescent mice received two-weeks of nicotine water (Adol Nic) or plain water (Adol Water). In adulthood, mice underwent three morphine tests: conditioned place preference (CPP), locomotor sensitization, and two-bottle choice. *Ex vivo* ventral tegmental area (VTA) brain slices were assessed via patch clamp for GABA and dopamine (DA) neuron morphine responses. Finally, VTA GABA neurons were chemogenetically inhibited during morphine CPP.

**Results:** In adulthood, Adol Nic mice had greater morphine CPP, more morphine locomotor sensitization, and more choice-based oral morphine consumption vs. Adol Water mice. In contrast, adult mice given nicotine vs. water had similar morphine CPP. Patch clamp analysis of VTA neurons from adult Adol Water mice showed canonical cell-type responses to bath-applied morphine: fewer action potentials in GABA neurons and more in DA neurons. Paradoxically, VTA GABA and DA neurons from adult Adol Nic mice did not show these morphine responses. In support of a causal relationship between GABA neuron firing and reward behavior, chemogenetic inhibition of VTA GABA neurons in Adol Water mice during pairing increased morphine CPP. In contrast, inhibition of VTA GABA neurons in Adol Nic mice brought morphine CPP down to control levels.

**Conclusions:** These data reveal an electrophysiological mechanism by which adolescent nicotine intake promotes morphine reward later in life, showing that adolescent nicotine exposure alters reward circuitry well into adulthood.

## INTRODUCTION

Opioid use disorder (OUD) claims thousands of lives (1,2) and costs the U.S. billions of dollars (3). A key, yet poorly understood, risk factor for OUD is the use of nicotine (4), the main addictive component in tobacco and e-cigarettes (5). After years of decline, cigarette smoking is rising again, driven by e-cigarettes (6). Adolescent nicotine has surged (7,8), with nearly one-third of teenagers reporting use (7,9). Epidemiological and animal studies link prior nicotine to increased intake of other addictive drugs, including opioids (10–21). However, the neural mechanisms underlying this relationship remain unknown.

Adolescence is a critical neurodevelopmental period, marked by major reorganization of limbic regions involved in reward processing (22). Both clinical and preclinical data show that adolescent drug exposure induces persistent behavioral changes (22–24), and adolescent nicotine exposure enhances drug reinforcement in adulthood (19,20,25–27). While adolescent nicotine exposure induces long-term adaptations in rodent brain function (19,28,29), how these neuroadaptations contribute to increased morphine reward remains unclear.

The ventral tegmental area (VTA) is a heterogenous midbrain region essential for addiction-related behaviors (30–37). Morphine infusion into the rodent VTA supports self-administration and place conditioning (38,39). The VTA consists primarily of dopamine (DA) neurons but also contains GABAergic and glutamatergic neurons (40,41). DA projections to the nucleus accumbens form the primary brain “reward” pathway (42–55), but GABA and glutamate neurons also regulate reward (56,57) often by modulating DA activity. VTA GABA neurons inhibit neighboring DA neurons and thus regulate DA release. Morphine’s acute rewarding effects partly stem from inhibiting VTA GABA neurons, leading to DA neuron disinhibition (58–60). Morphine binds to mu opioid receptors, predominantly on GABA neurons in the VTA, causing hyperpolarization and subsequent DA neuron disinhibition. Recent studies show that VTA GABA neurons participate in reward learning (61,62) and are disrupted by drugs of abuse (63–65). Chronic nicotine exposure in rodents dysregulates VTA GABAergic signaling (19,64,66–68), influencing DA signaling and behavioral responses to morphine (66,69). However, no studies have directly linked VTA GABA neuron activity changes to morphine reward, particularly following adolescent nicotine.

We postulate that adolescent nicotine enhances adult morphine reward by altering VTA GABA neuron responses. To test this, we used behavioral, electrophysiological, and chemogenetic approaches in mice. We found that adolescent nicotine increased adult morphine conditioned place preference (CPP), two-bottle choice morphine consumption, and morphine locomotor sensitization. Correspondingly, VTA GABA neurons exhibited a blunted response to morphine, diverging from the morphine-induced decrease seen in non-nicotine mice. VTA DA neurons also showed a blunted response to morphine, contrasting with the expected morphine-induced increase in DA activity (70,71). Finally, chemogenetic manipulations established a causal link between adolescent nicotine-induced VTA GABA dysfunction and adult enhanced morphine reward. These results indicate that reducing VTA GABA neuron firing during adult morphine exposure prevents the heightened morphine preference seen after adolescent nicotine.

## MATERIALS AND METHODS

### Animals and Ethics

Male and female wildtype C57BL/6J mice (Jackson Laboratory [JAX], #000664) and male heterozygous VGAT-Cre mice (homozygous VGAT-ires-Cre knock-in, JAX#028862 crossed with C57BL/6J mice) were used. Procedures were carried out in compliance with guidelines specified by the University of Pennsylvania Institutional Animal Care and Use Committee. This study adhered to guidelines for data management (randomization, blinding, etc. (72,73) (**Supplementary Materials [Supp. Mat.]**). For adolescent nicotine exposure, mice were deliberately group-housed despite this preventing individual intake measurements, as social isolation during adolescence produces robust stress effects that can permanently alter addiction-related behaviors and interact with nicotine’s effects on reward circuitry (16,74). Where sample sizes permitted, data were analyzed for sex differences as detailed in **Supp. Table 1**.

### Drugs

Nicotine hydrogen tartrate salt (Glentham Life Sciences) was given as 5-8mg nicotine/kg/mouse/day (75–78). Morphine sulfate (Spectrum) was given/used as 13-20mg/kg/mouse for two-bottle choice (79,80), 10mg/kg IP for CPP and locomotor sensitization (81), and 10uM for electrophysiology (82,83)). Clozapine N-oxide (CNO) dihydrochloride was given as 3 mg/kg IP (Hello Bio; (84,85)) and 50μM for slice electrophysiology experiments. This CNO concentration for slice experiments, while relatively high, was verified to produce reliable inhibition of VTA GABA neurons. Meloxicam as 2 mg/kg SC (**Supp. Mat.**). Drug details and all other experimental resources are provided in **Supp. Table 2**.

### Behavioral Tests

Testing started at P70 for adolescent nicotine experiments or at P112 for adult nicotine experiments. P70 testing consisted of morphine CPP, two-bottle choice, and locomotor sensitization (86–91). P112 testing consisted of morphine CPP (**Supp. Mat.**). Statistical analyses used the Sidak multiple comparisons test to control Type 1 error while maintaining reasonable statistical power.

### Stereotaxic Surgery

VGAT-Cre heterozygous mice received bilateral VTA infusion of a virus that expressed either an inhibitory DREADD (AAV5-DIO-hM4Di-mCherry, Addgene 44362-AAV5) or a control reporter protein AAV5-DIO-mCherry (Addgene 50459-AAV5) (92–95). For electrophysiology, mice received bilateral infusions of AAV9-CAG-FLEX-tdTomato (Addgene 28306-AAV9; **Supp. Mat.**).

### Ex Vivo Electrophysiology

Slice recordings were performed as previously described (19,21,96–98) (**Supp. Mat.**) using a Leica VT 1200S vibratome (Leica Microsystems). For DREADD validation experiments, VGAT-Cre mice were exposed to nicotine or water in adolescence, followed by bilateral VTA injection of cre-dependent hM4Di at P49. After three weeks of expression, VTA GABA neuron firing was recorded before and after bath-application of 50 μM CNO **(Supp. Mat.)**.

### Tissue collection, Immunohistochemistry (IHC), and Microscopy

Slide-mounted and free-floating IHC were performed as previously described (99,100) (**Supp. Mat.).**

### Data Analysis

**Supp. Table 1** provides statistical approaches, data structures, and results. Each figure legend provides a statistical analysis summary (101). Electrophysiological data was analyzed as previously published (21,96,97,102).

## RESULTS

### Adolescent, but not adult, oral nicotine exposure promotes morphine reward in adulthood

Past work showed male mice given nicotine in adolescence had higher morphine reward learning in adulthood (20). To test how adolescent nicotine influenced adult morphine CPP in both sexes, mice were exposed to nicotine or plain water for two weeks (P28-42; **Fig. 1A**). Daily fluid consumption was similar between Adol Nic and Adol Water groups, though within the adolescent nicotine group, intake was dose-dependent (**Fig. 1B-D**). Weight gain was comparable across groups (**Supp. Fig. 1**). After four weeks drug-free, adult (P70) Adol Nic mice of both sexes showed two-fold greater preference for the morphine-paired chamber vs. Adol Water mice (**Fig. 1E-F**), despite similar locomotion (**Fig. 1G**). There was no sex effect (**Supp. Fig. 2**), indicating adolescent nicotine enhances adult morphine reward independent of locomotion or sex.

**Figure 1:**
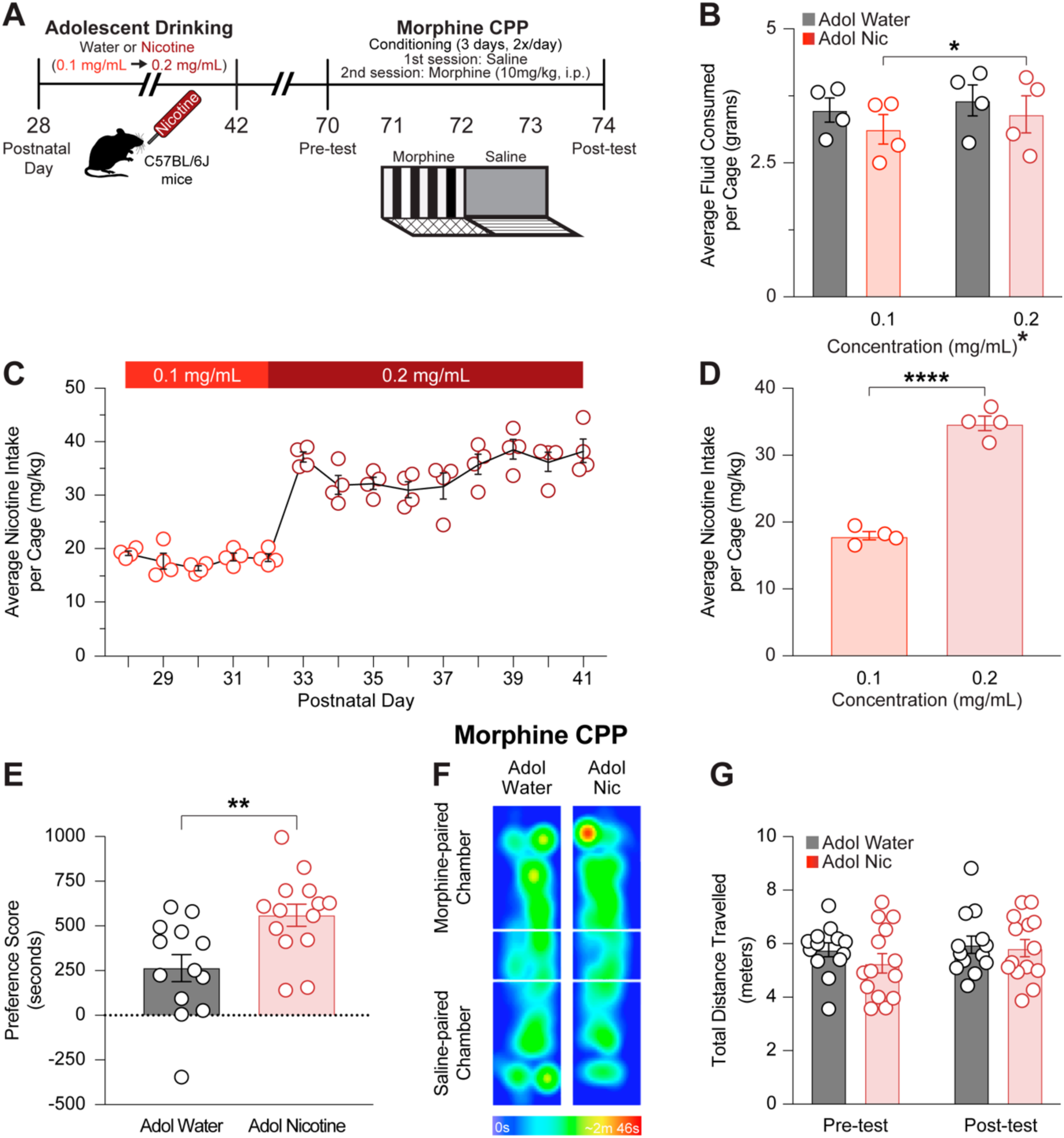
Nicotine exposure in the drinking water during adolescence promotes morphine conditioned place preference (CPP) in adulthood. **(A)** Experimental design. C57BL/6J male (n=12) and female (n=15) group-housed mice received 24-hour, continuous access to nicotine dissolved in their drinking water or plain water only from postnatal day (P) 28-42. CPP testing began at P70, consisting of 5 days (8 sessions, 30 min Post-test (Day 5). **(B)** Daily fluid consumption was similar between adolescent nicotine-exposed (red) and water (black) groups at both concentrations (two-way RM ANOVA: F(1, 6) = 0.6077, p=0.4653), with a main effect of concentration (F(1, 6) = 13.65, p=0.0102). Nicotine-exposed mice increased consumption at 0.2 mg/mL (Sidak’s post hoc, p=0.0386). Groups: Nicotine (n=14; 6 male, 8 female; 4 cages), Water (n=13; 6 male, 7 female; 4 cages). **(C)** Average nicotine intake per cage (n=4 cages; 2 male, 2 female cages). Nicotine concentration increased from 0.1 mg/mL to 0.2 mg/mL on Day 6. **(D)** Nicotine consumption increased at 0.2 mg/mL compared to 0.1 mg/mL (two-tailed paired t-test, ****p<0.0001). **(E)** At P70, morphine CPP testing (10 mg/kg, i.p. for three conditioning days) revealed 2-fold higher preference for the morphine-paired chamber in nicotine-exposed versus water mice (two-tailed unpaired t-test, **p=0.0052). **(F)** Representative heatmaps of animal positions during the Post-test. **(G)** Locomotion was similar between groups during Pre-test and Post-test (two-way RM ANOVA: F(1, 25) = 0.4021, p=0.5318). Data presented as mean ± SEM, *p<0.05, **p<0.01, ****p<0.0001.

To test if nicotine’s effect on morphine CPP was age-dependent, adult mice (P70-84) received nicotine or plain water for two weeks, followed by morphine CPP at P112 (**Fig. 2A**). Unlike adolescents, Adult Nic and Adult Water mice showed similar preference for the morphine-paired chamber (**Fig. 2B**) and locomotion (**Fig. 2C**). Thus, adolescence — but not adult — nicotine enhances morphine reward.

**Figure 2:**
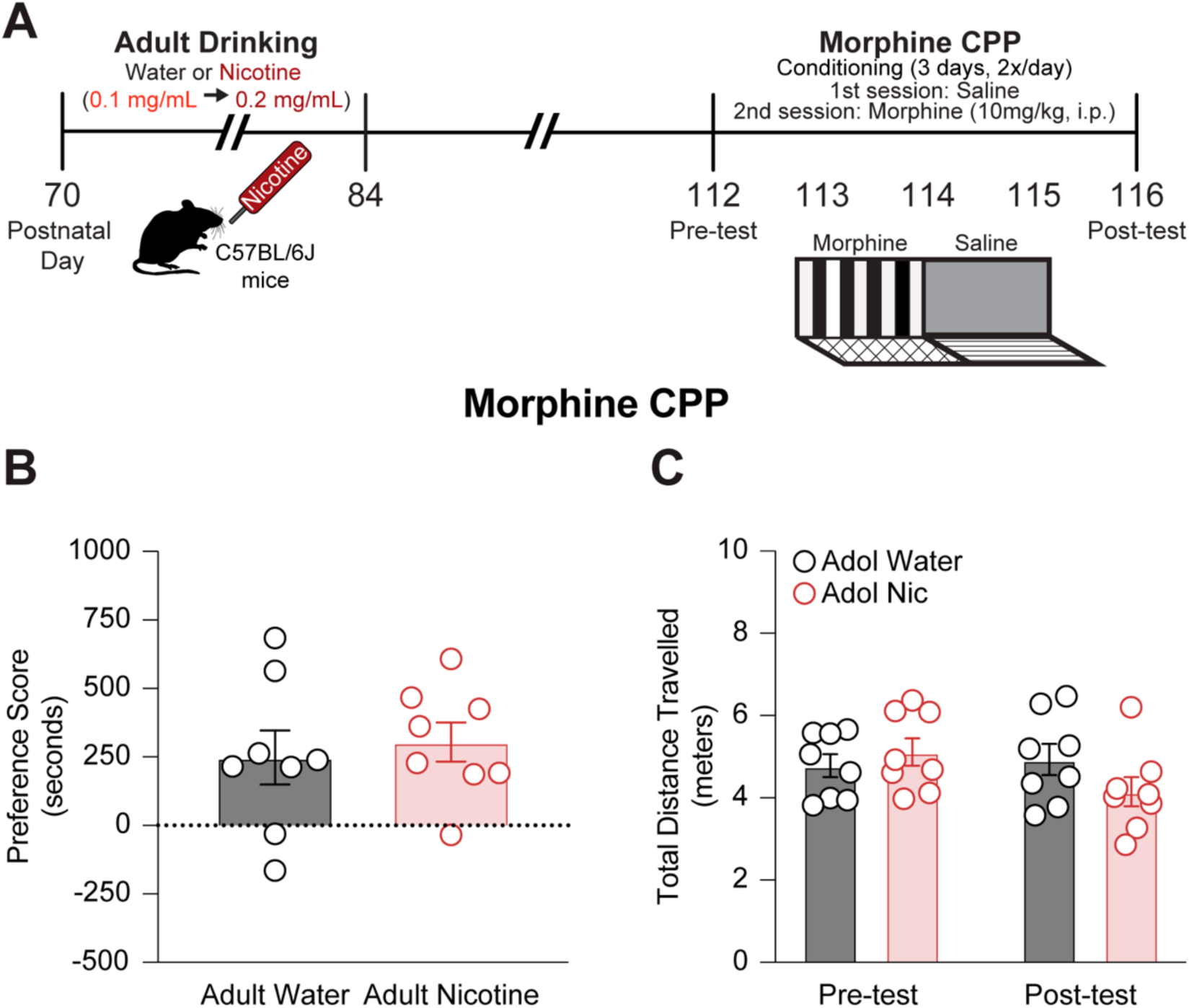
Nicotine exposure in the drinking water during adulthood does not impact adult morphine conditioned place preference (CPP). **(A)** Experimental design. C57BL/6J mice (n=16; 8 male, 8 female) received 24-hour continuous access to either nicotine solution or plain water from postnatal day (P) 70-84. CPP testing began at P112. **(B)** Preference for the morphine-paired chamber was similar between adult nicotine-exposed and water groups (two-tailed unpaired t-test, p=0.6509). **(C)** Locomotion did not differ between groups during Pre-test and Post-test (two-way RM ANOVA: F(1, 14) = 3.875, p=0.0691). Data presented as mean ± SEM.

Beyond CPP, we tested voluntary morphine consumption in adulthood using a two-bottle choice assay. Following adolescent nicotine or water exposure, adult (P67) mice were single-housed and acclimated to two saccharin bottles (**Fig. 3A**), showing no baseline preference (**Fig. 3B**). Beginning at P70, one bottle contained morphine dissolved in saccharin, increasing in concentration from 0.1 to 0.2mg/mL after nine days. Adol Nic mice consumed more morphine on 11 days (**Fig. 3C**) and sharply increased intake at the higher dose, while Adol Water mice maintained stable intake. Across all days and doses, Adol Nic mice drank more morphine than Adol Water mice (**Fig. 3D-E**). Notably, dose-dependent morphine intake occurred only among Adol Nic mice (**Fig. 3E**). These data suggest adolescent nicotine increases both reward learning and voluntary opioid consumption in a sex-independent manner (**Supp. Fig. 3**).

**Figure 3:**
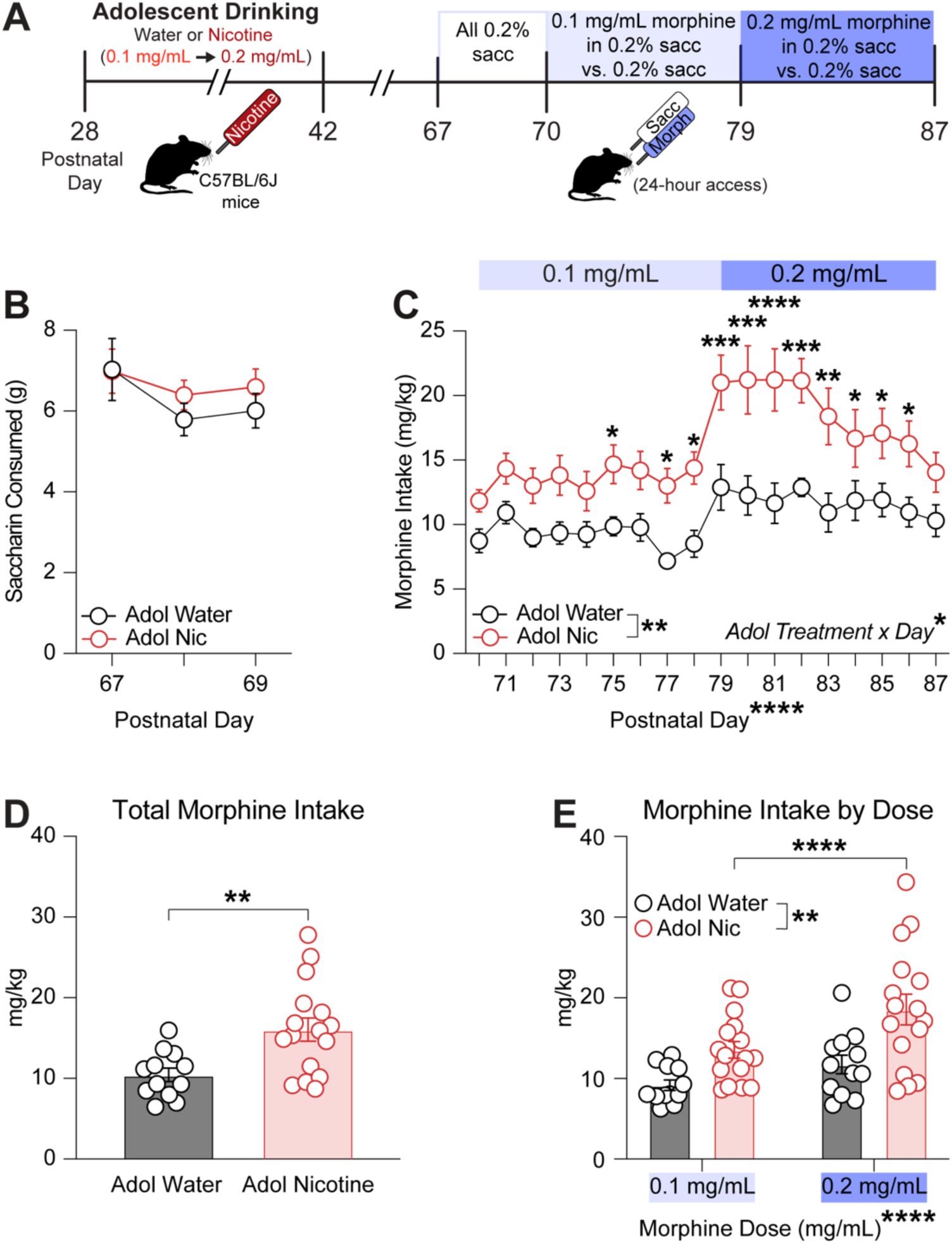
Adolescent nicotine increases consumption of morphine in adulthood in a two-bottle choice drinking paradigm. **(A)** Experimental design. C57BL/6J mice received 24-hour continuous access to nicotine solution or plain water from postnatal day (P) 28-42. At P67, mice were single-housed with access to two bottles containing 0.2% saccharin solution. After 3 baseline days, one saccharin bottle was replaced with morphine (0.1 mg/mL in 0.2% saccharin, Days 1-9; increased to 0.2 mg/mL, Days 10-18). **(B)** Saccharin intake was similar between groups during baseline days (two-way RM ANOVA: F(2, 52) = 0.5379, p=0.5872). Groups: Water (n=12; 7 male, 5 female), Nicotine (n=16; 7 male, 9 female). **(C)** Adolescent nicotine mice consumed more morphine than water controls (two-way RM ANOVA: interaction F(17, 442) = 1.991, p=0.0108; main effect of Adolescent Treatment F(1, 26)=9.615, p=0.0046; main effect of Day F(17, 442) = 10.69, p<0.0001). Sidak’s post hoc revealed higher consumption in nicotine group on Days 6, 8-17 (*p<0.05, **p<0.01, ***p<0.001, ****p<0.0001). **(D)** Total morphine intake, collapsed across dose and day, was higher in the nicotine group (two-tailed unpaired t-test, p=0.0046). **(E)** Nicotine-exposed mice showed increased consumption at higher dose (two-way RM ANOVA: interaction F(1, 26) = 2.848, p=0.1035; main effect of Adolescent Treatment F(1, 26)=9.615, p=0.0046; main effect of Dose F(1, 26)=27.24, p<0.0001). Within the nicotine group, consumption increased at 0.2 mg/mL versus 0.1 mg/mL (Sidak’s post hoc, p<0.0001). Data presented as mean ± SEM, *p<0.05, **p<0.01, ***p<0.001, ****p<0.0001.

### Adolescent nicotine exposure enhances morphine sensitization

Behavioral sensitization often co-occurs with changes in mesolimbic DA transmission (37). Thus, we tested whether adolescent nicotine affects adult locomotor sensitization to morphine. Mice of both sexes were exposed to nicotine or plain water during adolescence. Four weeks later, mice underwent an 11-day morphine sensitization paradigm (**Fig. 4A**). Both groups exhibited sensitization, with greater locomotion on the last vs. first day. However, despite similar movement between groups on the first day, Adol Nic mice traveled a greater distance on the last day compared to Adol Water mice (**Fig. 4B-C**), confirming enhanced morphine sensitization. These findings suggest adolescent nicotine exposure increases morphine addiction-related behaviors in adulthood.

**Figure 4:**
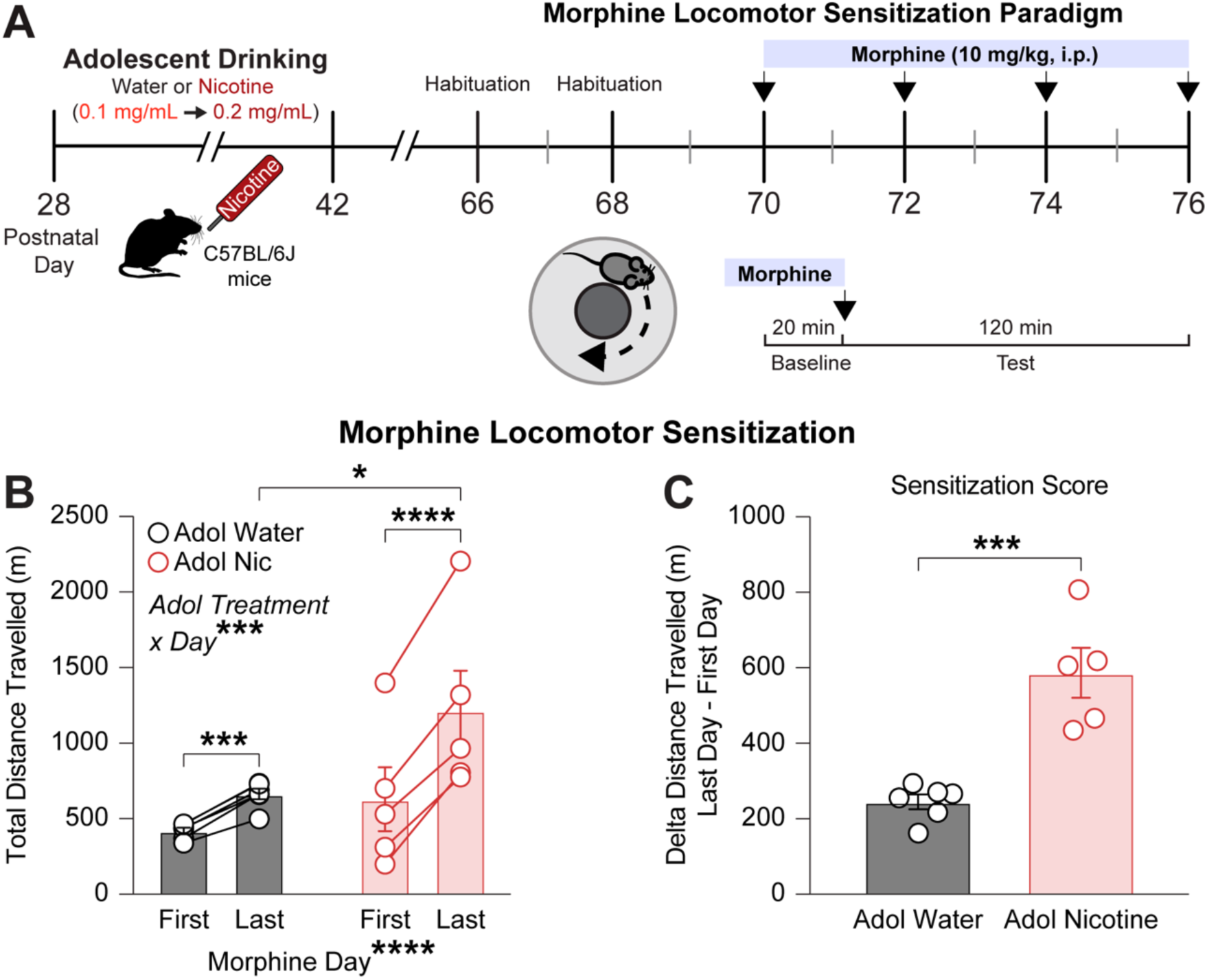
Adolescent nicotine exposure enhances morphine sensitization in adult mice. **(A)** Experimental timeline: C57BL/6J mice received either nicotine-containing or plain drinking water from postnatal day (P) 28-42. At P67, mice underwent morphine sensitization testing over 11 days, consisting of 6 sessions (2 saline habituation, 4 morphine) conducted every other day. Each session included a 20-minute baseline period followed by 2-hour post-injection monitoring. **(B)** Two-way repeated measures ANOVA revealed significant Adolescent Treatment x Day interaction (F(1,9) = 29.08, p=0.0004) and a main effect of Time (F(1,9) = 171.4, p<0.0001). Both adolescent water-treated and adolescent nicotine-treated mice expressed morphine sensitization, as there was more morphine-induced locomotion on the last relative to the first testing day (Sidak’s post hoc, p=0.0006 and p<0.0001, respectively). In addition, there was a significant difference in Total Distance Travelled between Adolescent Water and Adolescent Nicotine on the Last Day (Sidak’s post hoc, p=0.0427). Group composition: Adolescent Water: n=6 (5 males, 1 female); Adolescent Nicotine: n=5 (2 males, 3 females). **(C)** Sensitization magnitude (last day minus first day locomotion) was significantly higher in adolescent nicotine-treated mice compared to water controls (two-tailed unpaired t-test, p=0.0004). Data shown as mean ± SEM. *p<0.05, ***p<0.001, ****p<0.0001.

### Adolescent nicotine exposure alters VTA GABA neuron firing rate in response to morphine

Chronic drug exposure alters VTA GABA transmission (19,103–105), and adolescent nicotine induces lasting changes in VTA GABA neuron function (19). To test adolescent nicotine’s impact on morphine responses *ex vivo*, VGAT-Cre mice received adolescent nicotine or plain water, followed by AAV9-CAG-FLEX-tdTomato infusions at P56 to label VTA VGAT+ neurons. Patch clamp recordings at P70 identified VTA GABA neurons (**Fig. 5A-B, Supp. Fig. 4A-C**). Baseline firing rates were similar across groups (**Fig. 5C**). After bath-application of 10 μM morphine (71,82,83), VTA GABA neurons in Adol Water mice had dramatically reduced firing (58,59,106) **(Fig. 5D-E)**. However, VTA GABA neurons from Adol Nic mice showed little or no inhibition, with some even increasing firing (**Fig. 5F**). Over time, Adol Nic neurons fired more than Adol Water neurons at multiple time points (**Fig. 5G**), independent of sex (**Supp. Fig. 5**). Since baseline rates were unaffected (**Fig. 5C**), this effect is not due to a floor effect. Thus, in contrast to adolescent water, adolescent nicotine exposure prevents morphine-induced inhibition of VTA GABA neurons in adulthood.

**Figure 5:**
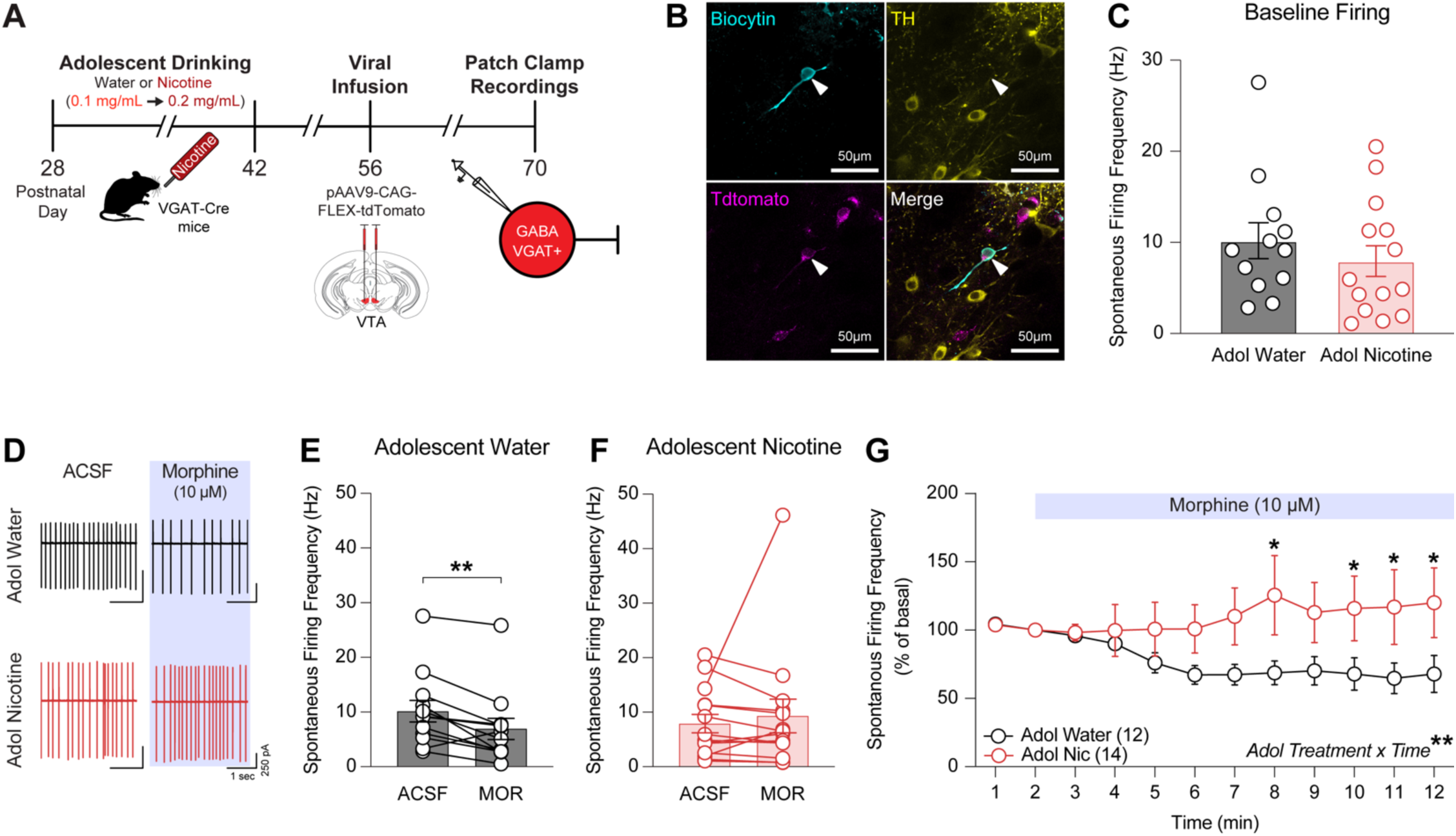
Prior adolescent nicotine exposure alters morphine-induced inhibition of ventral tegmental area (VTA) GABA neuron firing in adult mice. **(A)** Experimental timeline: VGAT-Cre mice received nicotine-containing or plain drinking water from postnatal day (P) 28-42. At P56, mice received bilateral infusions of Cre-dependent tdTomato virus to label GABA neurons. At P70, spontaneous firing rates of VTA GABA neurons were recorded in cell-attached configuration before and after morphine (10 μM) bath application. **(B)** Representative biocytin-filled neuron visualized with streptavidin labeling. Scale bar: 50 μm. **(C)** Baseline spontaneous firing frequencies did not differ between groups prior to morphine application (two-tailed unpaired t-test, p=0.3939). Measurements taken during second minute of baseline recording. Group composition: adolescent water (n=12 cells from 6 mice; 2 males, 4 females), adolescent nicotine (n=14 cells from 10 mice; 7 males, 3 females). Data shown as mean ± SEM. **(D)** Representative traces from adolescent water-exposed (black) and nicotine-exposed (red) mice. **(E)** VTA GABA neurons from adolescent water-exposed mice showed significant reduction in firing rate between baseline (ACSF, minute 2) and morphine treatment (minute 12) firing rates between baseline and morphine treatment (Wilcoxon matched-pairs signed rank test, p=0.3575). **(G)** Time course of normalized firing rates following morphine application. Two-way repeated measures ANOVA revealed significant group differences over time (F(11,264) = 2.709, p=0.0025). Sidak’s post-hoc analysis showed significant differences between groups at minutes 8, 10, 11, and 12 (p<0.05). Data shown as mean ± SEM. *p<0.05, **p<0.01.

### Adolescent nicotine exposure changes VTA DA neuron firing rate in response to morphine

VTA GABA cells regulate local VTA DA neurons and project to other brain regions. To investigate the circuit-level impact of adolescent nicotine, we examined VTA DA neuron responses to morphine in adulthood. VGAT-Cre mice received adolescent nicotine or plain water, followed by AAV9-CAG-FLEX-tdTomato VTA infusions at P56. Patch clamp recording at P70 identified non-fluorescent, TH+ DA neurons (**Fig. 6A-B**). Baseline DA neuron firing was similar between groups (**Fig. 6C**). Among Adol Water mice, morphine increased DA neuron firing (70,71) (**Fig. 6D-E**). However, among Adol Nic mice, morphine did not change DA firing (**Fig. 6D-E**). Over time, Adol Water DA neurons fire more than Adol Nic neurons at several time points (**Fig. 6F**), with no baseline differences (**Fig. 6C**), ruling out a ceiling effect. These data suggest adolescent nicotine exposure prevents morphine-induced excitation of VTA DA neurons in adulthood, implicating local VTA GABA interneurons in this altered response.

**Figure 6:**
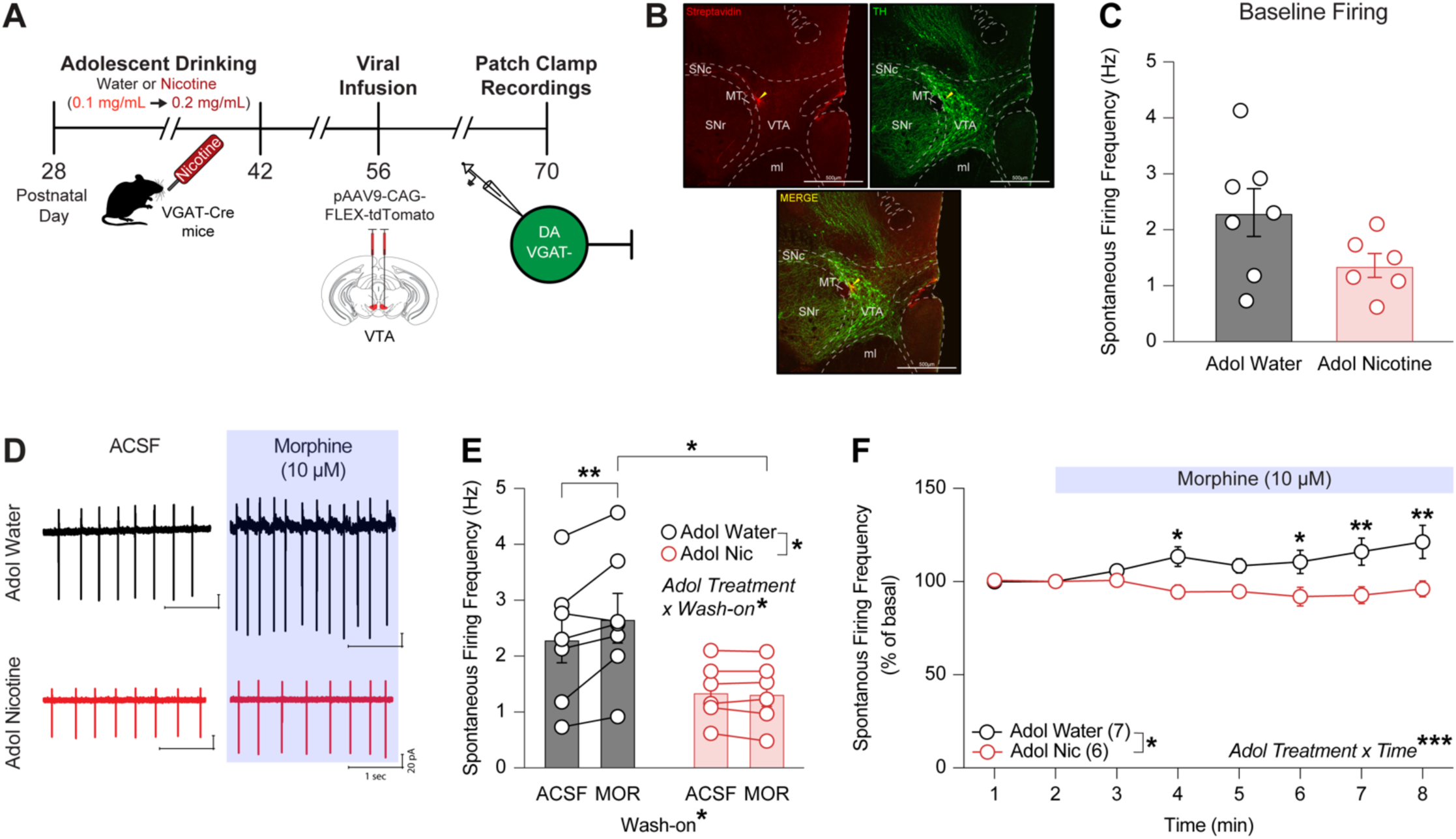
Prior adolescent nicotine exposure attenuates morphine-induced excitation of ventral tegmental area (VTA) dopamine neurons in adult mice. **(A)** Experimental timeline: VGAT-Cre mice received nicotine-containing or plain drinking water from postnatal day (P) 28-42. At P56, mice received bilateral infusions of Cre-dependent tdTomato virus to label GABA neurons. At P70, spontaneous firing rates of VTA dopamine neurons (identified as VGAT-negative/non-fluorescent cells) were recorded in cell-attached configuration before and after morphine (10 μM) bath application. **(B)** Representative biocytin-filled neuron visualized with streptavidin labeling. Scale bar: 500 μm. **(C)** Baseline spontaneous firing frequencies did not differ between groups prior to morphine application (two-tailed unpaired t-test, p=0.0878). Measurements taken during second minute of baseline recording. Group composition: adolescent water (n=7 cells from 5 mice; 2 males, 3 females), adolescent nicotine (n=6 cells from 4 mice; 1 male, 3 females). Data shown as mean ± SEM. **(D)** Representative traces from adolescent water-exposed (black) and nicotine-exposed (red) mice. **(E)** Two-way repeated measures ANOVA revealed significant Adolescent Treatment × Wash-on interaction on (F(1,11) = 5.623, p=0.0371). VTA dopamine neurons from adolescent water-exposed mice showed increased firing between baseline (minute 2) and morphine treatment (minute 8) (Sidak’s post-hoc, p=0.0063). Firing rates significantly differed between groups at minute 8 (Sidak’s post-hoc, p=0.0326). **(F)** Time course of normalized firing rates following morphine application. Two-way repeated measures ANOVA revealed significant effects of time (F(7,77) = 4.325, p=0.0004) and Adolescent Treatment (F(1,11) = 8.203, p=0.0154). Sidak’s post-hoc analysis showed significant differences between groups at minutes 4, 6 (p<0.05), and 7, 8 (p<0.01). Data shown as mean ± SEM. *p<0.05, **p<0.01, ***p<0.001.

### Paradoxical firing of VTA GABA neurons is necessary for adolescent nicotine-induced enhanced morphine CPP

To determine if altered VTA GABA neuron activity contributes to enhanced morphine CPP, we tested if chemogenetic inhibition of VTA GABA neurons prevents this effect. VGAT-Cre mice received adolescent nicotine or water, followed by AAV5-hsyn-DIO-hM4Di-mCherry or control AAV5-hsyn-DIO-mCherry infusions into the VTA (**Fig. 7A-B, Supp. Table 3**). Mice were divided into 4 groups (**Fig. 7C**) and underwent morphine CPP.

**Figure 7:**
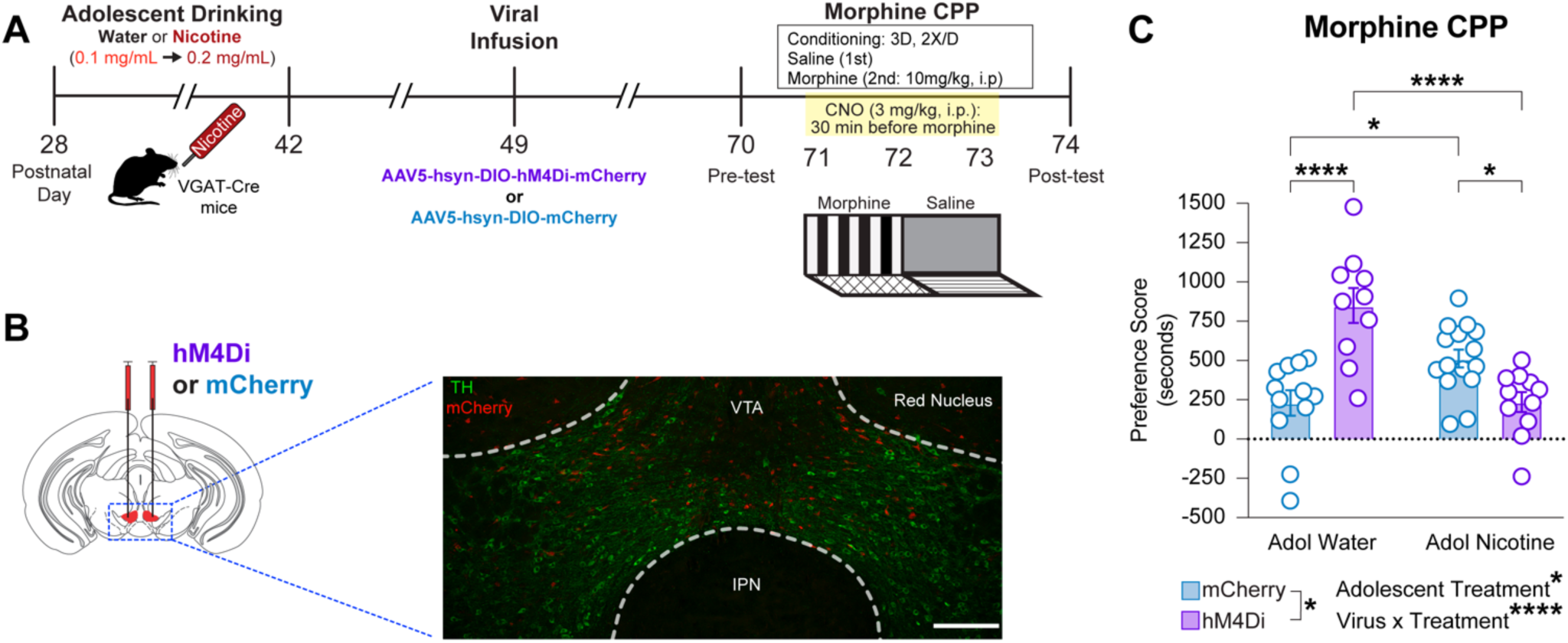
Chemogenetic inhibition of ventral tegmental area (VTA) GABA neurons prevents enhanced morphine conditioned place preference (CPP) in adult mice with adolescent nicotine exposure. **(A)** Experimental timeline: VGAT-Cre mice received nicotine-containing or plain drinking water from postnatal day (P) 28-42. At P49, mice received bilateral VTA infusions of Cre-dependent virus expressing either inhibitory DREADD (hM4Di-mCherry) or control fluorophore (mCherry). Morphine CPP testing began 3 weeks post-surgery. **(B)** Representative image showing bilateral VTA viral expression. Immunofluorescence for mCherry (red) shows viral expression, and tyrosine hydroxylase (TH, green) defines VTA boundaries. Scale bar: 200 μm. Mice with virus expression outside VTA boundaries were excluded from analysis. **(C)** Morphine CPP (10 mg/kg, i.p.) was assessed in mice expressing hM4Di-mCherry or mCherry control. CNO (3 mg/kg, i.p.) was administered 30 minutes before each morphine conditioning session to activate DREADDs. Two-way ANOVA revealed significant Virus × Adolescent Treatment interaction (F(1,44) = 33.51, p<0.0001), and main effects of Adolescent Treatment (F(1,44) = 4.575, p=0.038) and Virus (F(1,44) = 4.887, p=0.0323). Sidak’s post-hoc analysis showed significant differences in morphine chamber preference between: Adolescent water-mCherry vs nicotine-mCherry (p=0.0171); Adolescent water-mCherry vs water-hM4Di (p<0.0001); Adolescent water-hM4Di vs nicotine-hM4Di (p<0.0001); Adolescent nicotine-mCherry vs nicotine-hM4Di (p=0.0232). Group composition: Adolescent water-mCherry: n=12 (6 males, 6 females); Adolescent nicotine-mCherry: n=15 (11 males, 4 females). Adolescent water-hM4Di: n=10 (3 males, 7 females); Adolescent nicotine-hM4Di: n=11 (7 males, 4 females). Data shown as mean ± SEM. *p<0.05, ****p<0.0001.

hM4Di, when activated by CNO, suppresses neuronal activity via Gi-coupled signaling. Since Adol Nic VTA GABA neurons resist morphine inhibition (which also occurs through Gi-coupled signaling), we confirmed that hM4Di remained functional in these neurons (**Supp. Fig. 6C-E**). Adol Nic-mCherry mice expressed greater preference for the morphine-paired chamber than Adol Water-mCherry mice (**Fig. 7C**), consistent with **Fig. 1D**. However, chemogenetic inhibition of VTA GABA neurons (Adol Nic-hM4Di + CNO) blocked this enhanced CPP. This suggests abnormal VTA GABA neuron firing is necessary for adolescent nicotine-induced enhancement of morphine reward. Interestingly, Adol Water-hM4Di mice had a higher CPP score than Adol Water-mCherry and Adol Nic-hM4Di mice. Since morphine reward is typically mediated by VTA GABA neuron inhibition and thus DA disinhibition, we propose a model where adolescent nicotine prevents normal GABA suppression, thereby driving heightened morphine CPP. Conversely, inhibiting VTA GABA neurons in drug-naïve mice enhances opioid reward (107). These results indicate that adolescent nicotine disrupts normal morphine reward circuitry, engaging a novel reward mechanism.

## DISCUSSION

Nicotine use often begins in adolescence, a uniquely vulnerable period during which drug exposure has long-lasting neural and behavioral ramifications (24). Clinical and preclinical data reveal that nicotine use enhances the rewarding effects of multiple drugs of abuse, including alcohol, cocaine, and opioids (4,14,19, 20). Here we demonstrate that adolescent nicotine exposure fundamentally alters how morphine engages reward circuits in adulthood. Mice exposed to nicotine during adolescence show enhanced morphine reward in adulthood across multiple behavioral measures: increased conditioned place preference, greater voluntary consumption, and heightened locomotor sensitization. This effect is specific to adolescent exposure, as nicotine given in adulthood did not alter subsequent morphine reward. Most strikingly, we identify a novel mechanism whereby adolescent nicotine exposure transforms how VTA GABA neurons respond to morphine, leading to enhanced reward through sustained — rather than reduced — GABA activity.

The canonical model of morphine reward involves inhibition of VTA GABA neurons, which then disinhibits DA neurons to drive reward. Previous work has also found that the inhibition of GABA neurons alone is reinforcing (108), and our control data robustly support this established mechanism. VTA GABA cells from adolescent water-exposed mice show the expected decrease in firing in response to morphine (58) while their DA neurons show increased firing (70,71). Adolescent nicotine exposure fundamentally rewires this circuit, as demonstrated by our comprehensive recordings from both cell types. VTA GABA neurons from nicotine-exposed adolescent mice paradoxically maintain their firing in response to morphine, while their DA neurons consequently fail to show the typical increase in firing. This coordinated transformation in both cell types provides compelling evidence for local circuit reorganization as a key mechanism underlying how sustained GABA activity results in heightened reward. The maintained GABA firing directly prevents the normal morphine-induced increase in DA neuron activity, a relationship we causally validated through chemogenetic manipulations: inhibiting VTA GABA neurons during morphine conditioning specifically prevents the heightened preference seen in nicotine-exposed mice.

The insensitivity of VTA GABA neurons to morphine following adolescent nicotine exposure drives this paradoxical enhancement of reward through multiple mechanisms. While GABA neurons typically increase their firing in response to morphine, our recordings reveal a complete loss of this canonical response, suggesting fundamental alterations in opioid signaling. This GABA neuron insensitivity likely disrupts the balance between tonic and phasic dopamine release, where altered GABA activity could reduce tonic DA firing while enhancing the signal-to-noise ratio of phasic DA bursts. Our dopamine recordings indicate that local VTA GABA interneurons are particularly affected, pointing to adaptations in local circuit connectivity that transform how GABA activity influences DA release. Additionally, distinct populations of VTA GABA neurons may show different adaptations - long-range GABAergic projections to the nucleus accumbens influence associative learning, while rostral VTA GABAergic projections to dorsal raphe nucleus modulate morphine reward (109,110). The differential sensitivity of these GABA populations to adolescent nicotine exposure may orchestrate the enhanced reward phenotype.

Beyond local VTA reorganization, adolescent nicotine exposure likely also induces broader neuroadaptations across the reward circuitry that could amplify these effects. These include alterations in nucleus accumbens synaptic plasticity and receptor expression, changes in prefrontal cortical control over VTA-NAc signaling affecting reward seeking, adaptations in stress-responsive circuits including the extended amygdala, and modifications to cholinergic signaling in VTA-projecting regions. These circuit-wide changes likely work synergistically with our observed VTA adaptations to enhance morphine’s rewarding properties.

At the cellular level, several mechanisms could explain why VTA GABA neurons from nicotine-exposed adolescent mice resist morphine-induced inhibition. Morphine typically inhibits GABA neurons through mu opioid receptor-mediated activation of inhibitory signaling cascades. One possibility is that adolescent nicotine exposure alters this signaling cascade. Alternatively, compensatory circuit-level adaptations may emerge, such as through VTA glutamate neurons that express mu opioid receptors (111) and could excite neighboring GABA neurons. The diversity of VTA GABA neurons likely contributes to these complex adaptations, as these neurons differ in their synaptic targets and molecular composition (40). While canonical markers of interneuron subtypes from forebrain regions do not directly map onto the VTA (40), and calcium-binding proteins often co-express in both GABA and DA neurons (112), this heterogeneity may explain both our observed cell-to-cell variability and the novel reward mechanism we report.

Within our behavioral results, it is important to consider that nicotine exposure during adolescence could alter taste perception in ways that influence our two-bottle choice drinking results. While we attempted to control for bitter taste by dissolving morphine in saccharin, chronic nicotine exposure can modify taste processing in rodents (113–115). However, two aspects of our data suggest that altered taste sensitivity alone cannot explain our results. First, the dose-dependent increase in consumption specifically in nicotine-exposed adolescent mice suggests an interaction with morphine’s rewarding properties rather than just reduced bitter taste detection. Second, our CPP and sensitization findings, where taste is not a factor, support that adolescent nicotine exposure enhances morphine’s rewarding properties through mechanisms beyond taste alone.

These findings challenge the canonical view of VTA GABA neurons in reward processing. While optogenetic activation of VTA GABA neurons typically produces aversion and interrupts reward consumption (116,117), and their inhibition is rewarding (107), these neurons also encode reward expectation (61,118). Our results reveal that adolescent nicotine exposure creates an alternative pathway to reward that operates through sustained, rather than reduced, GABA activity. This fundamentally changes our understanding of how early nicotine exposure affects reward circuitry and may help explain why adolescent nicotine use increases vulnerability to opioid addiction.

## ACKNOWLEDGMENTS

This work was supported by National Institute on Drug Abuse Grants F31 DA053111 (PI: REW) and R37 DA053296 (PI: JAD) as well as National Institute on Alcohol Abuse and Alcoholism R01 AA026267 (PI: JAD). This work was also funded by National Institute of Mental Health R01 MH129970 (PI: AJ Eisch), National Institute of Diabetes and Digestive and Kidney R01 DK135871 (PI: SA Zedric), National Institute of Neurological Disorders and Stroke R01 NS088555 (PI: AM Stowe) and a 2021 NASA Hero grant 80NSSC21K0814 (PI: S Yun). Moreover, the Eisch Lab is supported by philanthropic funds from an anonymous donor and the Dani Lab is supported by a generous award from the Chernowitz Medical Research Foundation. We also acknowledge the generous intellectual support of the members of the Dani and Eisch Labs.

## DECLARATION OF COMPETING INTEREST

The authors declare no conflicts of interest.

## SUPPLEMENTARY MATERIAL

### SUPPLEMENTARY MATERIALS AND METHODS

#### Animals

Male and female wildtype C57BL/6J mice, postnatal day 14 (P14; The Jackson Laboratory [JAX], #000664) arrived with the dam in the temperature- and humidity-controlled vivarium facility at the University of Pennsylvania. They were allowed to acclimate to a reverse light/dark cycle (lights off 10 AM, lights on 10 PM) for one week prior to weaning at P21 (4 mice/cage). At the same time as weaning, mice were ear punched for identification purposes. These mice were used for morphine conditioned place preference (CPP), morphine two-bottle choice, and morphine locomotor sensitization experiments. In addition, male homozygous VGAT-Cre mice (VGAT-ires-Cre knock-in (C57BL/6J), JAX#028862) were crossed with female wildtype C57BL/6J mice resulting in heterozygous VGAT-Cre mice bred in-house at the University of Pennsylvania. Only F1 were used. Mice were genotyped by PCR using genomic DNA and previously published primers. Heterozygous VGAT-Cre mice were used for electrophysiological recordings and *in vivo* chemogenetic manipulations. All home cages were provided with environmental enrichment and *ad libitum* food and water. Mice were group-housed at all times, unless otherwise indicated, in order to avoid isolation stress. All procedures were carried out in compliance with guidelines specified by the Institutional Animal Care and Use Committee at the University of Pennsylvania.

#### Drugs

Nicotine hydrogen tartrate salt (Glentham Life Sciences, #GL9693-1G) was dissolved in filtered water (nicotine concentration: 0.1mg/mL or 0.2 mg/mL) for adolescent or adult drinking (based on drinking ∼4 ml of water/mouse/day) for a final dose range of 5 to 8 mg nicotine/kg/mouse/day (1–4). Morphine sulfate (Spectrum Chemical and Laboratory Products, Catalogue No: M1167) was dissolved in filtered water (morphine concentration: 0.1 mg/ml or 0.2 mg/ml) for two-bottle choice drinking experiments for a final dose range of 13 to 20 mg/kg/mouse (5,6), for CPP and locomotor sensitization in sterile saline (0.9% sodium chloride) (Hospira, Inc., NDC: 0409-1918-32; final morphine dose via IP injection: 10 mg/kg (7)), and in ACSF for electrophysiological recordings (morphine concentration: 10uM (8,9)). All concentrations for nicotine and morphine are provided as freebase equivalents. Clozapine N-oxide (CNO) dihydrochloride (Hello Bio, Catalogue No: HB6149) was dissolved in sterile saline for IP injection (CNO dose 3 mg/kg, (10,11)). Meloxicam (Loxicom Norbrook Company, Catalogue No: 2091-90A; 5 mg/mL) was dissolved in sterile saline fresh for each SC injection and injected at a final concentration of 2 mg/kg.

#### Nicotine Treatment

Adolescent nicotine treatment occurred from P28-42 to correspond with the midpoint of adolescence (12). Group-housed mice were given 24-hour, continuous access to nicotine dissolved in their drinking water (0.1 mg/mL on Days 1-5; 0.2 mg/mL on Days 6-14) or water only. Adult nicotine treatment occurred from P70-84 to correspond with when the adolescent treated group was assessed for adult behaviors. For this adult nicotine treatment, group-housed mice again received either nicotine in water (0.1 mg/mL on Days 1-5; 0.2 mg/mL on Days 6-14) or plain water. While preventing the collection of individual intake data and subsequent correlation with measures of morphine behavior, group-housing mice during the adolescent or adult drinking period was a deliberate choice for several reasons: a) social isolation during adolescence produces robust stress effects that can permanently alter addiction-related behaviors (13), b) adolescents are particularly vulnerable to isolation stress (14,15), c) prior work shows stress interacts with nicotine’s effects on reward circuitry (16), and d) single-housing would have introduced a significant confound, making it impossible to distinguish between the effects of nicotine exposure vs. isolation stress. The experimental choice to group-house mice prioritized maintaining normal social development during the critical period of adolescence to ensure the findings specifically reflect nicotine’s effects.

#### Behavioral Tests

Behavior testing started at P70 for the adolescent nicotine experiment or at P112 for the adult nicotine experiment. The P70 behavior testing consisted of morphine CPP, morphine two-bottle choice, and morphine locomotor sensitization, performed as previously described (17–22). The P112 behavior testing consisted of morphine CPP. Brief descriptions of the behavioral tests are provided below. Male and female mice were used for all behavioral tests and all testing was completed during the dark cycle. For CPP and locomotor sensitization, mice were acclimated to the behavior room and daily sham needle intraperitoneal insertion for 5 days prior to the start of testing.

##### Morphine CPP

###### CPP Chamber

Mice underwent CPP pairing (conditioning) and testing in an unbiased, 3-compartment apparatus (22,23). Each house-made CPP apparatus had overall dimensions of 61 cm (length) x 15 cm (width) x 33 cm (height). Each apparatus consisted of three chambers: 2 equal-sized outer chambers (pairing compartments, each 24.5 cm (length) x 15 cm (width) x 33 cm (height)) joined by a smaller center compartment (12 cm (length) x 15 cm (width) x 33 cm (height)). The smaller center compartment could be isolated from the outer chambers by temporary partitions. Each of these 3 compartments offered a distinct textural and colored environment. The gray pairing compartment consisted of gray walls and hard plastic floors with grooves in a striped pattern; the striped pairing compartment consisted of black and white striped walls and hard plastic floors with grooves in a criss-cross pattern; and the middle compartment consisted of white walls with metal bar flooring. Lighting in the room was dimmed to be 7 lux in all 3 CPP compartments. No differences in time spent in the middle compartment were observed for groups tested (data not shown). The design of this box resulted in an unbiased conditioning apparatus, and a counterbalanced compartment assignment was used for morphine and saline pairings (20).

###### CPP Pairing and Testing

Mice underwent pairing and testing in morphine CPP with a 5-day paradigm (17,18) beginning at P70 (for the adolescent nicotine experiment) or P112 (for the adult nicotine experiment). Briefly, mice were first acclimated to the behavior room for 5 days, and received daily sham needle intraperitoneal insertion each day. Then at the start of the experiment (on Day 1, Pretest, 3-5 PM), mice were placed in the middle CPP compartment. At the start of the session, the temporary partitions were raised, and mice had full access to the CPP box for 30 min. Time spent in each compartment was recorded by the ANY-maze (Stoelting Co.) video tracking software. Mice with greater than 60% (18 min) of time spent in any compartment were excluded to ensure subjects did not have an initial bias in the apparatus. Days 2-4 consisted of 30 minute conditioning sessions, both a saline session and a morphine session on each day. On these pairing days, the CPP box was divided into 3 compartments (striped, middle, and gray compartments) with temporary partitions, enabling mice to be confined to their assigned compartment. The first session (11 am-3 pm) was always the saline (sterile, bacteriostatic 0.9% saline, i.p.)-paired context (conditioned stimulus, CS-) and the second session (5 pm-9 pm) was always the morphine (10 mg/kg, i.p.).-paired context, (CS+). Mice were randomly assigned to either a gray or striped CS+ compartment, with the alternate context serving as the CS-compartment. An equal number of mice were conditioned with morphine (10 mg/kg, i.p.) in the gray and striped compartments. Morphine was always administered during the second pairing session to increase the time between the last morphine dose and the next conditioning session. This limits the possible confound of residual morphine in the system influencing the next conditioning session. The timing of the conditioning sessions was equally framed around the timing of the Pretest and Posttest (which were each done 3-5PM) to avoid any confounds related to testing at the same time as only one of the conditioning sessions. Day 5 consisted of the Posttest (3-5 PM). Mice were again placed in the middle compartment and at the start of the session, the temporary partitions were raised; mice received full access to the CPP box for 30 minutes. As with the Pretest, the time spent in each compartment was recorded by ANY-maze video tracking.

###### CPP Scoring

The CPP or “preference” score was calculated as Post-test (time spent on morphine-paired side – time spent on saline-paired side) – Pre-test (time spent on morphine-paired side – time spent on saline-paired side).

###### Chemogenetics and CPP

For the chemogenetic CPP experiment (see below), the CPP procedure was the same as described above with the addition of an injection of CNO (3 mg/kg, i.p.) administered to all groups 30 minutes prior to the start of each morphine conditioning session.

##### Morphine Two-Bottle Choice

A separate group of mice were single-housed with environmental enrichment (i.e. nestlet and shepherd shack) starting at P67, 3 days before beginning the morphine two-bottle choice drinking experiment at P70. They were acclimated to two bottle access during this three-day habituation period by providing two 50-ml black sipper tubes containing 0.2% (w/v) saccharin (Sigma-Aldrich, St. Louis, MO) in the home cage, placed on opposite sides of the cage. Days were defined as beginning at 11 am and ending at 11 am the subsequent day. Data from the three-day habituation period showed there was no preference for the bottle on either side of the cage (data not shown for side preference). Following the three-day saccharin habituation period, the experiment began at Day 1 when one bottle was replaced with 0.1 mg/mL morphine dissolved in 0.2% saccharin. The other bottle remained filled with 0.2% saccharin. In half of the cages, the morphine bottle was placed on the left side and in the other half of the cages, the morphine bottle was placed on the right side. This continued from Days 1-9. From Days 10-18, the dose of morphine was increased to 0.2 mg/mL dissolved in 0.2% saccharin and the other bottle remained filled with 0.2% saccharin. Since there was no preference for the bottle on either side of the cage, morphine and saccharin positions within each cage did not change throughout the course of the experiment. Both solutions were provided 24 hours a day, 7 days a week. All bottle measurements were adjusted for leakage (i.e. bottle weight minus solution loss from bottles placed in an empty cage), with final bottle weights recorded in grams. Mice were weighed at the start of each drinking day. No other fluid was provided throughout the duration of the drinking experiment. Morphine consumption was analyzed across each 24-hour period accounting for body weight (mg/kg). Preference for the morphine solution relative to saccharin was also analyzed across each 24-hour period (expressed as a percentage, where >50% indicates morphine preference). Analysis of preference ratios (morphine solution/total fluid consumed) revealed that while neither group showed absolute preference for morphine over saccharin, adolescent nicotine-exposed mice showed significantly higher preference scores (main effect of Adolescent Treatment, p=0.0198) and maintained higher consumption across doses.

##### Morphine Locomotor Sensitization

Morphine locomotor sensitization was generally performed as previously described (19). Beginning at P70, mice were first acclimated to the behavior room for 5 days and received daily sham needle intraperitoneal insertion each day. Then mice underwent two chamber habituation days with one day of rest in between. For each habituation day, they were placed into the one foot in diameter circular apparatus and locomotor activity was measured (ANY-maze) for 20 minutes to establish a within-subject baseline, and then received an injection of saline and locomotor activity was recorded for an additional 120 minutes. Locomotor activity from the second day of saline treatment was considered the baseline for statistical analyses. After these saline habituation days, the mice then underwent four days of morphine locomotor sensitization (with one day of rest in between each day of sensitization). For sensitization, mice received 20 min in the chamber mice then all received morphine (10 mg/kg, i.p.). Development of behavioral sensitization to morphine was defined as a significant increase in locomotor activity between the first and last (fourth) day of morphine treatment.

#### Stereotaxic Viral Infusions

VGAT-Cre heterozygous mice received bilateral VTA infusions of a virus that either expressed an inhibitory chemogenetic protein (AAV5-DIO-hM4Di-mCherry, Addgene 44362-AAV5) or a control reporter protein AAV5-DIO-mCherry (Addgene 50459-AAV5) following previously published work (24–27). In brief, mice were deeply anesthetized under isoflurane and placed into a stereotaxic apparatus. Bilateral infusions of an adeno-associated virus expressing AAV5-DIO-hM4Di-mCherry (Addgene 44362-AAV5, titer: 1×10^11^) or a control virus (AAV5-DIO-mCherry, Addgene 50459-AAV5, titer: 1×10^11^) were delivered to the VTA. For both viruses, the titers were 1×10^11^, the volume infused was 150 nL/side, the rate was 30 nL/min, and the delivery was via 33 gauge Hamilton syringes (Hamilton, Reno, NV). For electrophysiology experiments, mice received bilateral infusions of AAV9-CAG-FLEX-tdTomato (Addgene 28306-AAV9). VTA stereotaxic coordinates were: A/P: −3.25 mm, M/L: +/− 0.5mm, D/V: −4.55mm relative to Bregma. For all viruses, the volume infused was 150 nL/side, the rate was 30 nL/min, the delivery was via 33 gauge Hamilton syringes (Hamilton, Reno, NV), and needles were kept in place an additional 10 minutes to ensure diffusion. Mice were randomly assigned to experimental and control groups. Throughout the surgery, mice were periodically assessed for any reflex. Two to three weeks elapsed between surgery and electrophysiological or behavioral experiments to allow viral expression. Mice received the non-opioid analgesic, meloxicam, during surgery and every 12 hours for 2 days post-surgery to alleviate pain.

#### Ex vivo Electrophysiology

Slice recordings were performed as previously described (28–32). Briefly, P70 mice were anesthetized with isoflurane, decapitated, and the brain removed from the skull and placed in ice-cold oxygenated (95% O2/5% CO2) high-sucrose artificial cerebrospinal fluid (ACSF, in mM): 205.0 sucrose, 2.5 KCl, 21.4 NaHCO_3_, 1.2 NaH_2_PO_4_, 0.5 CaCl_2_, 7.5 MgCl_2_, 33.3 dextrose. Horizontal slices (260 μm) containing the VTA of mice were cut using a vibratome (Leica Microsystems, Model: Leica VT 1200S). Following cutting, slices were immediately transferred to oxygenated (95% O2/5% CO2), standard ACSF for 40 min at 32°C (in mM): 120 NaCl, 3.3 KCl, 25.0 NaHCO3, 1.2 NaH2PO4, 2.0 CaCl2, 1.0 MgCl2, 10.5 dextrose, 20.0 sucrose. Slices recovered at 32℃ and then at room temperature for at least 1 hour prior to recording. During recordings, slices were perfused with standard ACSF at 32℃ at a constant flow rate of 2-3 ml/min. Thin-walled borosilicate glass (1.12 mm ID, 1.5 mm OD; WPI) was used to make patch electrodes with resistances between 2.5-3.5 MΩ when filled with internal solution. A K-gluconate-based internal solution was used (in mM): 123.0 K+ gluconate, 8.0 NaCl, 2.0 Mg-ATP, 0.2 EGTA, 10.0 HEPES, 0.3 Tris-GTP, 0.4% biocytin (adjusted with KOH to pH 7.2-7.3 and 295mOsm with sucrose). The firing rates of VTA GABA neurons were recorded in cell-attached configuration in voltage clamp mode with the holding potential set at 0 mV (33).

To measure the effects of morphine (10 μM dissolved in ACSF, Spectrum) on action potential firing in GABA and DA neurons, stable firing was achieved for at least 2 minutes before bath application of morphine. We selected 10 μM of morphine because this concentration is commonly used in in vitro experiments (8) and the effect of morphine at this concentration can be abolished by the opioid receptor antagonist, naloxone (9). To measure the effects of CNO (50 μM dissolved in ACSF, Spectrum) (34) on action potential firing in GABA neurons, stable firing was also achieved for at least 2 minutes prior to bath application of 50 μM CNO, a relatively high dose. Pre-drug baseline and either morphine or CNO data points were collected for all cells. When possible, after cell-attached experiments, the patch configuration was converted to whole-cell and the h current (I_h_) was measured. Recorded neurons were fixed in 4% paraformaldehyde for post hoc immunohistochemistry against TH and streptavidin labeling of biocytin-filled cells. Analyses of electrophysiological data were performed using Clampfit and with custom-made protocols in Igor.

#### Tissue collection, Immunohistochemistry (IHC), and Microscopy

Slide-mounted and Free-floating IHC were performed as previously described (35,36).

##### Slide-mounted (SM) IHC on tissue from behaviorally tested mice

At the end of behavioral testing, mice were anesthetized with a combination of ketamine (80 mg/kg) and xylazine (10 mg/kg) and underwent intracardial exsanguination with cold 0.1M PBS (7 ml/min, 6 min) followed by transcardial perfusion with 4% paraformaldehyde in 0.1 M PBS (15 min) for analysis of viral expression. Brains were dissected and postfixed for 48 hours in 4% paraformaldehyde in 0.1 M PBS. After subsequent cryoprotection in 30% sucrose, brains were sectioned coronally on a freezing microtome (Leica), collecting 40 μm sections through the entire anterior-posterior length of the brain. Serial sets of sections were stored in 0.1% NaN_3_ in 1xPBS at 4°C until processing for SM IHC. For IHC, one series of sections was mounted on glass slides (Superfrost Plus, Fisher). Sections were processed for antigen retrieval (0.01 M citric acid, pH 6.0, 95°C, 15 min) and then after PBS rinses endogenous peroxidases were quenched for 30 minutes (0.3% H_2_O_2_ in 1 x PBS). Following rinses, nonspecific staining was prevented by incubating in blocking solution (3% normal donkey serum (NDS), in 0.1% Triton X-100 in 1xPBS) for 30 min. After blocking, sections were incubated in the following primary antibodies: rat anti-mCherry (1:3000, Invitrogen, #M11217) and rabbit anti-tyrosine hydroxylase (1:500, Abcam, #ab112) in 3% NDS and 0.1% Tween-20 in 1xPBS overnight for at least 16 hours. The next day, after PBS rinses, sections were incubated in the following secondary antibodies: biotinylated donkey anti-rat (Jackson Immunoresearch, #712-065-153) and donkey anti-rabbit Cy3 (Jackson Immunoresearch, #711-165-152) in 1.5% NDS and 1 x PBS for 1 hour. After rinses, sections were incubated in avidin-biotin complex (ABC Elite, Vector Laboratories, #PK-6100) for 60 minutes. After rinsing, staining was visualized using Fluorescein, Cyanine 3-labeled Tyramide Signal Amplification (TSA, Akoya Biosciences, #NEL701A001KT) for 10 minutes. After PBS rinses, sections were incubated in DAPI (Sigma, #236276) as a counterstain for 15 minutes. Finally, after rinsing, slides were dehydrated briefly in a series of alcohols: 70%, 95%, 100%, and then dipped in Citrasolv, coverslipped and left to dry overnight before imaging. Tissue was imaged at 10x and 20x on an epi-fluorescent microscope (Olympus, Center Valley, PA). Mice were excluded based on analysis of viral expression, i.e. any diffusion into structures neighboring the VTA. <u>FF IHC on slices used for patch clamping</u> was performed as previously described (36). Recorded slices were rinsed in 1 x PBS and processed for antigen retrieval (10 mM sodium citrate, pH 8.5, incubated for 30 minutes at 45℃). Slices were then rinsed again in PBS and incubated for 2 hours in 50 mM glycine at room temperature. Following another rinse, slices were incubated in a blocking solution for 3 hours at room temperature containing 5% bovine serum albumin (Sigma), 5% normal goat serum (Vector Laboratories) and 0.5% Triton-X (VWR) for 3 hours at room temperature. Afterwards, slices were incubated overnight for at least 16 hours at 4℃ in a primary antibody solution containing 1xPBS, 1% BSA, 1% NGS, 0.1% Triton-X, streptavidin Alexa Fluor 568 (1:2000; Thermo Fisher Scientific, #S11226) and rabbit anti-tyrosine hydroxylase (1:500, Abcam, #ab112). The following day, slices were again rinsed and then incubated for 6 hours at room temperature in a secondary antibody solution containing PBS, 1% BSA, 1% NGS, 0.1% Triton X and goat anti-rabbit Alexa Fluor 488 (1:500, Thermo Fisher Scientific, #A-11008). After a final rinse in 0.1M sodium phosphate buffer (pH 7.35), slices were mounted on glass slides (Superfrost Plus, Fisher) and coverslipped with fluorescent mounting medium (Fluoromount-G, Thermo Fisher Scientific).

#### Data Analysis (37)

Rout (Q=1%) was performed for all measures. Data are reported as mean +/− SEM. Prior to all statistical analyses, data assumptions (e.g. normal distribution) were verified. For normality testing, within-group distributions were assessed by the Shapiro-Wilk test, D’Agostino & Pearson test, and/or Kolmogorov-Smirnov test and visualized with Quantile-Quantile plots. All statistical approaches, data structures, and results are provided in **Supplementary Table 1** and statistical analysis summaries are provided in each figure legend. Analyses with two groups were performed using a paired or unpaired, two-tailed, t-test as appropriate. Analyses with more than two groups were performed using a one-way ANOVA and Sidak’s multiple comparison’s test, following the presence of a significant interaction or main effect. Post hoc significance is indicated on all plots by asterisks. Analyses with more than two variables were performed using a two-way ANOVA again followed by Sidak’s multiple comparison’s test in the event of a significant interaction or main effect. Sidak’s multiple comparison’s test was chosen because it provides better control of Type 1 error while maintaining reasonable statistical power. Repeated measures were used where appropriate and recorded in figure legends and **Supplementary Table 1**. All statistical analyses were performed using Prism (version 10.3.1). The significance level for all analyses was set at p < 0.05.

For electrophysiological data, analysis was performed as previously published (28,30,31,38). Specifically, voltage clamp traces containing spontaneous action potentials were imported into Clampfit (Molecular Devices), and each action potential was detected using their template-matching event detection protocol. Time stamps of the detected action potentials were exported into Excel (Microsoft) for further analysis. The data was converted into frequency measurements (Hz) in 60 second time bins, for 3 minutes prior to morphine application and then for 10 minutes during morphine wash-on. These data for each cell were normalized to the action potential frequency at minute 3, just prior to morphine application. Data was averaged within groups.

#### Randomization and Blinding (39,40)

Mice were not selected for any experiment or group based on previous observations or tests. Since mice were group-housed for all experiments, an entire cage received the same adolescent treatment and, if applicable, the same virus. Cages were selected arbitrarily to receive control or experimental adolescent treatments and viral injections. Individual mice within each cage were given ear punches for identification. Behavior chambers were numbered front to back, starting with box one, and the animal number determined which box the mouse experienced throughout testing. Care was taken that each box had equal numbers of control and experimental animals assigned to it over the course of an experiment (e.g. box 1 would not be assigned only to adolescent nicotine-treated or hM4Di mice). All behavioral experiments were controlled by computer systems and data were collected and analyzed in an automated and unbiased way. For the chemogenetic experiment, histological verifications always took place prior to analysis of behavioral data. For electrophysiology experiments, data were analyzed on a day separate from the recording and knowledge of the adolescent treatment was not known at the time of analysis.

### SUPPLEMENTARY FIGURES AND FIGURE LEGENDS

**Supplemental Figure 1:**
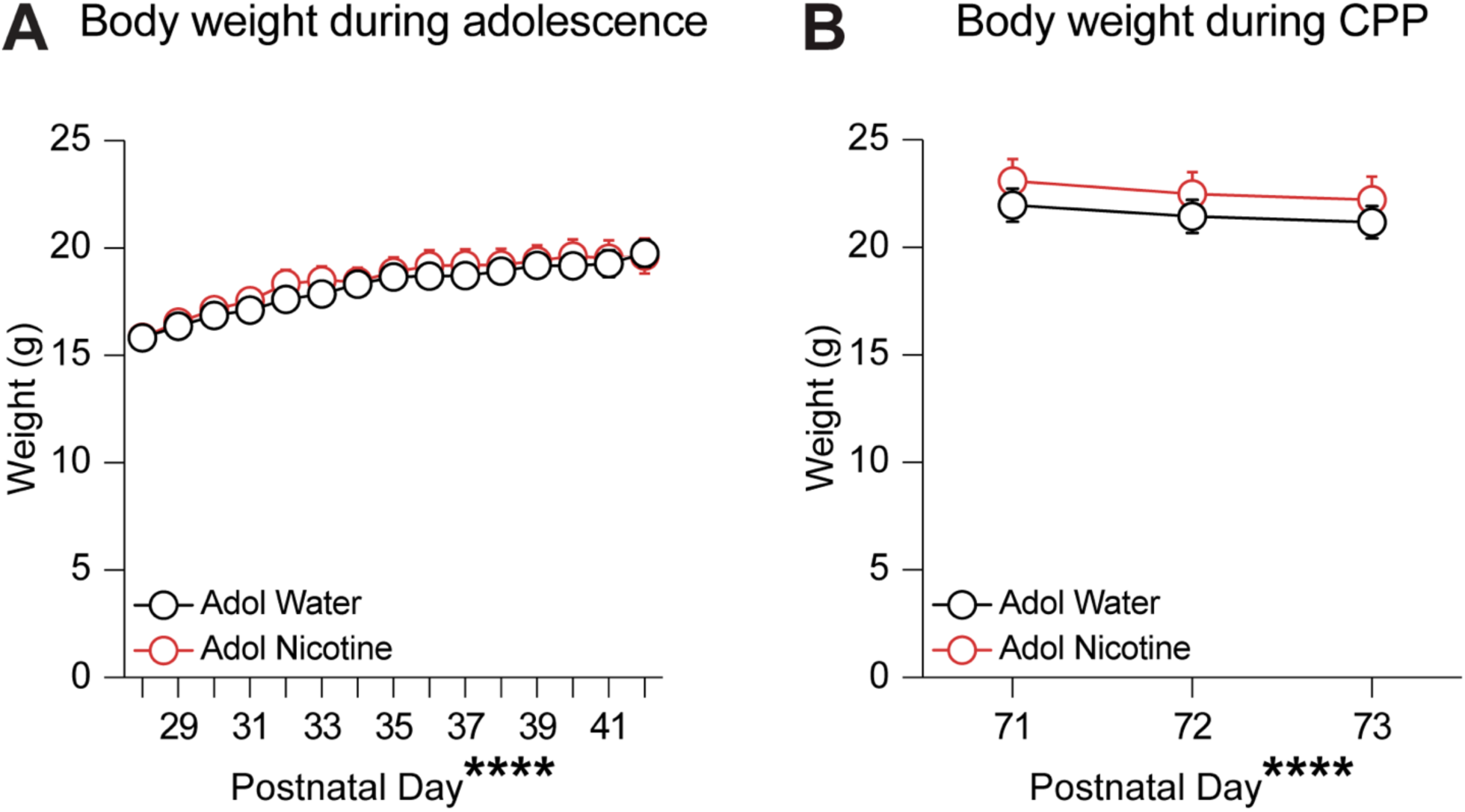
Body weight is similar throughout life in adolescent nicotine-treated and adolescent water-treated mice. **(A)** There was no significant difference in body weight between adolescent nicotine-treated (red) and adolescent water-treated (black) mice throughout the two-week adolescent drinking period (two-way RM ANOVA: F(14, 364) = 1.014, p=0.4383). There was a main effect of Day (F(14, 364) = 110.9, p<0.0001). Groups: Water (n=13; 6 male, 7 female), Nicotine (n=14; 6 male, 8 female). **(B)** Throughout the three morphine CPP conditioning days in adulthood, there were also no differences in body weight between groups (two-way RM ANOVA: F(2, 52) = 0.2472, p=0.7819). There was a main effect of Day (F(2, 52) = 79.21, p<0.0001). Data presented as mean ± SEM.

**Supplemental Figure 2:**
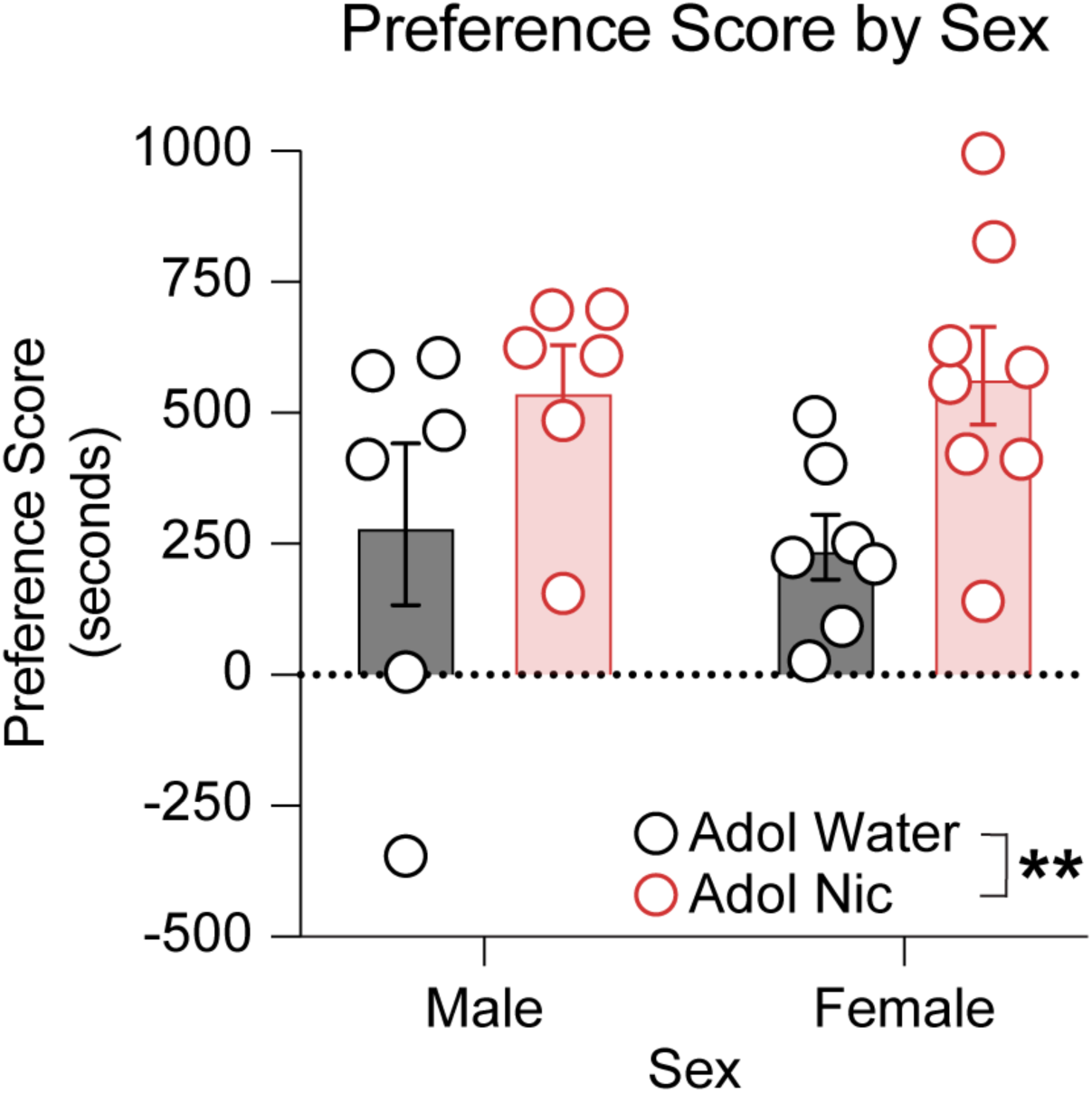
Nicotine exposure in the drinking water during adolescence promotes adult morphine CPP to a similar extent in male and female mice. There is no effect of sex on the adolescent nicotine-induced increase in adult morphine CPP (two-way ANOVA, F(1, 23) = 0.1188, p=0.7334). There is the main effect of adolescent treatment we previously report (F(1, 23) = 8.349, p=0.0083). Groups: Water (n=13; 6 male, 7 female), Nicotine (n=14; 6 male, 8 female). Data presented as mean ± SEM, **p<0.01.

**Supplemental Figure 3:**
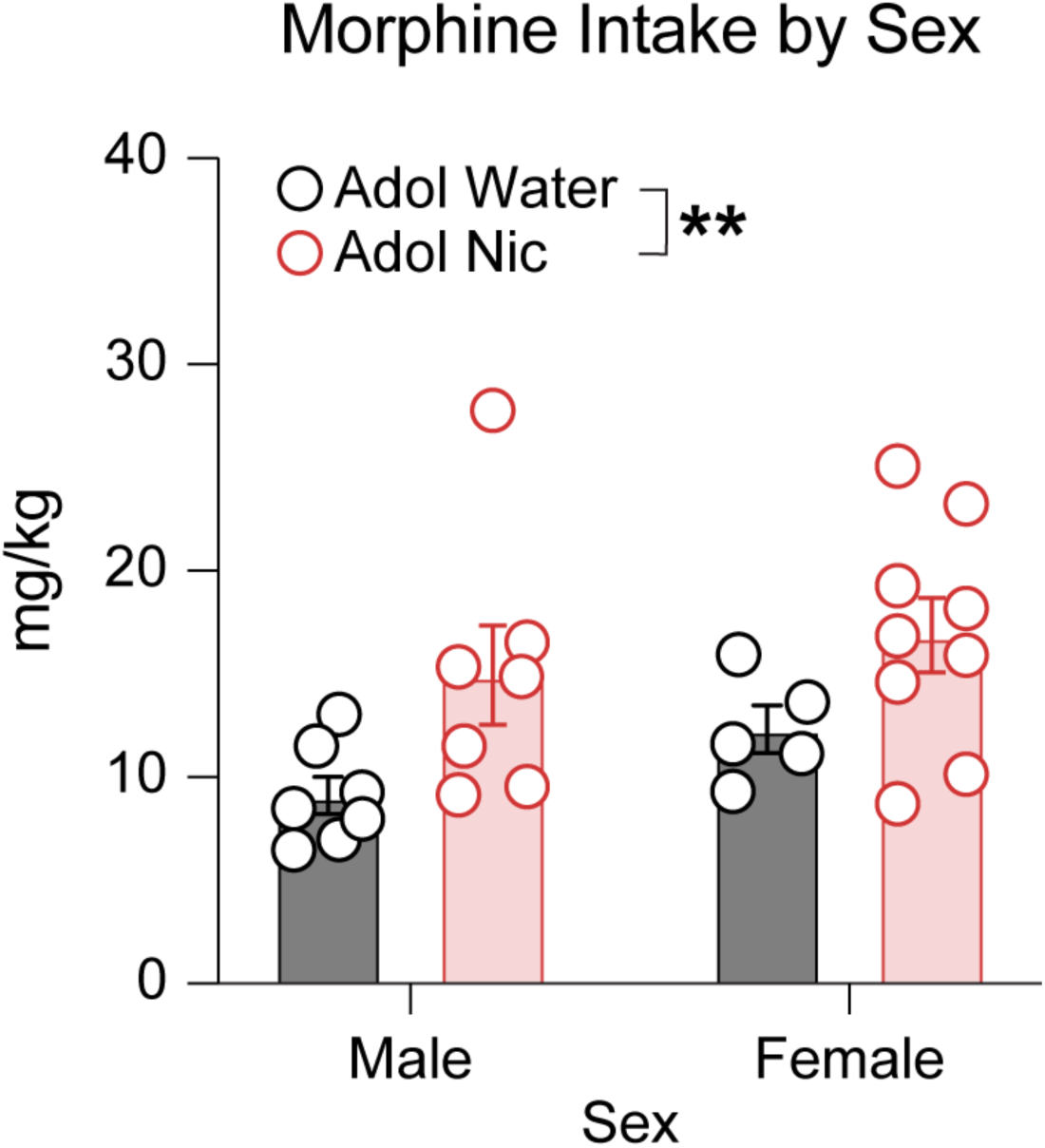
Nicotine exposure in the drinking water during adolescence increases morphine consumption in adulthood to a similar extent in male and female mice. There is no effect of sex on the adolescent nicotine-induced increase in adult morphine consumption (two-way ANOVA, F(1, 24) = 0.1254, p=0.7264). There is the main effect of adolescent treatment we previously report (F(1, 24) = 8.140, p=0.0088). Groups: Water (n=12; 7 male, 5 female), Nicotine (n=16; 7 male, 9 female). Data presented as mean ± SEM, **p<0.01.

**Supplemental Figure 4:**
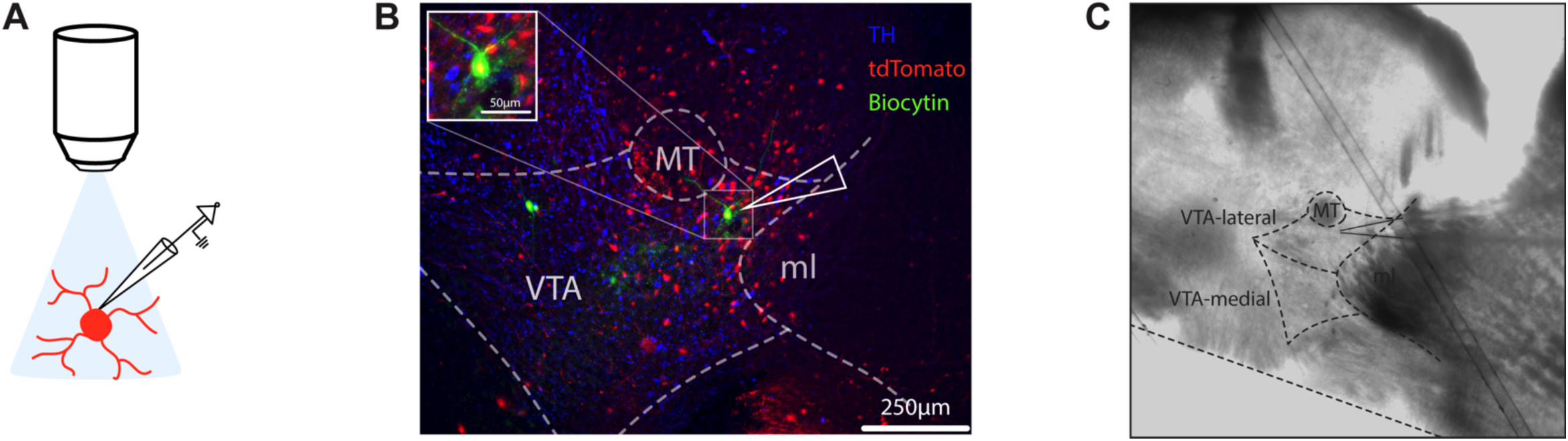
VTA GABA neuron identification and recording location. **(A)** VGAT-Cre mice received bilateral intra-VTA infusions of pAAV9-CAG-FLEX-tdTomato two weeks prior to recordings in order to genetically label GABA neurons for identification in slice. **(B)** TH expression delineating the borders of the VTA. Biocytin was added to the internal solution to enable post hoc verification of the identity and location of the recorded cell. Scale bar, 250 μm. **(C)** GABA neuron recording location in the lateral VTA, under DIC optics.

**Supplemental Figure 5:**
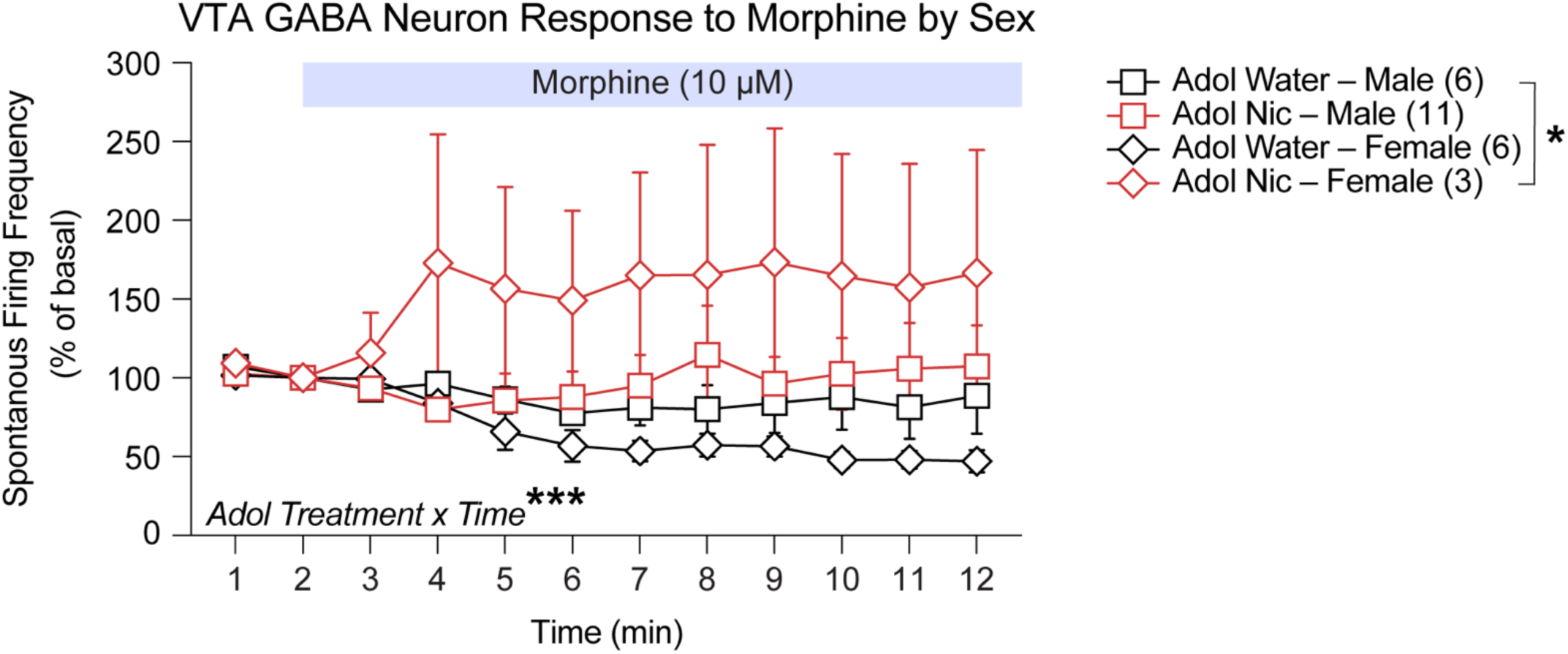
Independent of sex, VTA GABA neurons from adolescent nicotine-treated mice have a greater spontaneous firing rate in response to morphine compared to cells from adolescent water-treated mice. There is no effect of sex on the greater VTA GABA cell firing rate observed among adolescent nicotine-treated mice compared to adolescent water-treated mice (three-way RM ANOVA, F(11, 242) = 1.778, p=0.0585). There is the time x adolescent treatment interaction (F(11, 242) = 3.409, p=0.0002) we previously report as well as a main effect of adolescent treatment (F(1, 22) = 4.860, p=0.0382). Groups: Water-male (n=6 cells from 2 mice), Water-female (n=6 cells from 4 mice), Nicotine-male (n=11 cells from 7 mice), Nicotine-female (n=3 cells from 3 mice). Data presented as mean ± SEM, *p<0.05, ***p<0.001.

**Supplemental Figure 6:**
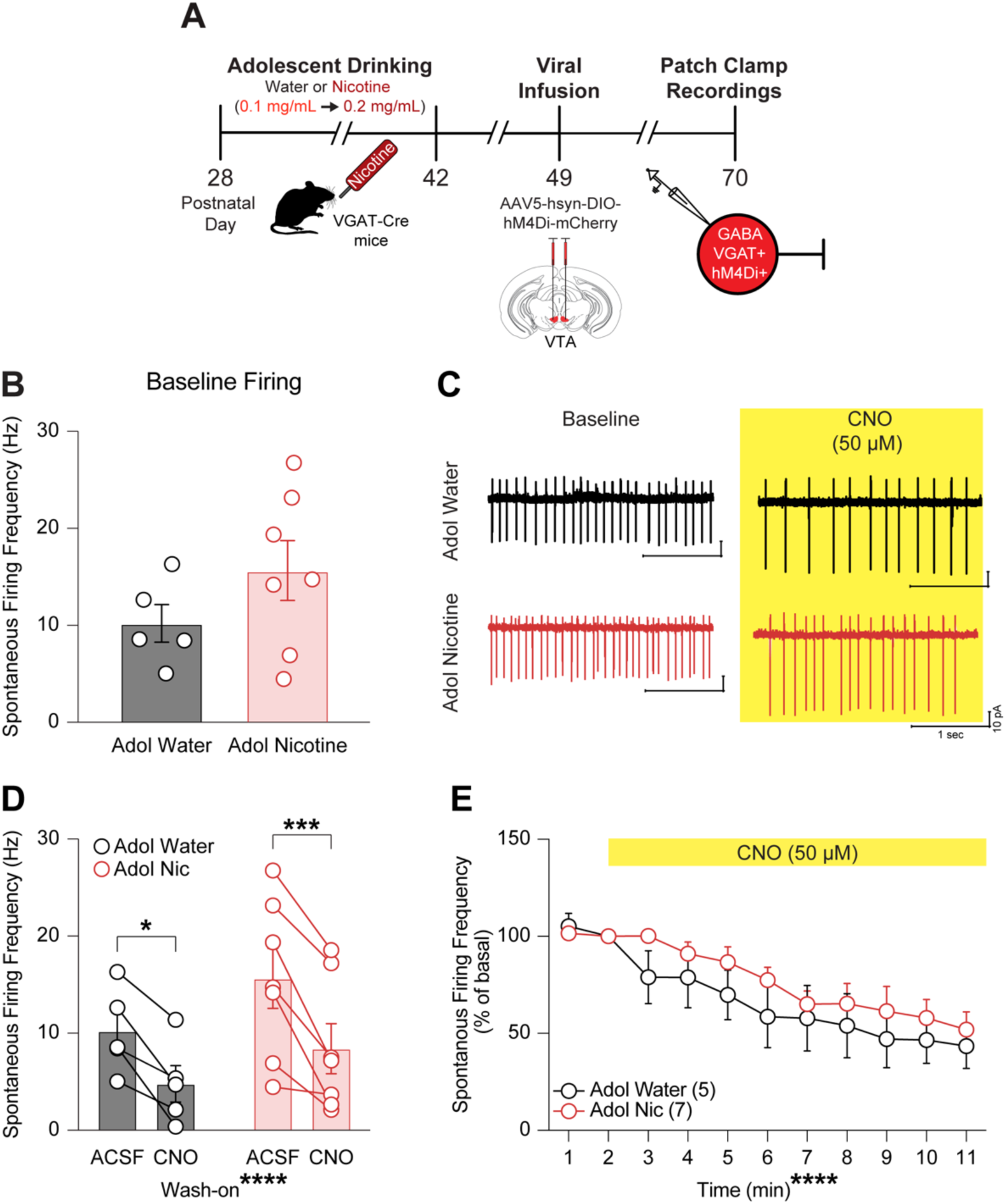
hM4Di inhibits VTA GABA neurons from both adolescent water-treated and adolescent nicotine-treated mice. **(A)** Experimental design. VGAT-Cre male and female group-housed mice received 24-hour, continuous access to nicotine dissolved in their drinking water or plain water from P28-42. At P49, mice receive bilateral VTA infusions of a cre-dependent viral construct expressing inhibitory (hM4Di) designer receptors exclusively activated by designer drugs (DREADD). Three weeks later, at P70, mCherry-expressing neurons were recorded in cell-attached mode to monitor spontaneous action potential firing before and after bath-application of CNO (50 μM). **(B)** In adulthood, there was no difference in baseline spontaneous firing frequency of cells from adolescent water-treated and adolescent nicotine-treated mice prior to CNO wash-on (two-tailed, unpaired t-test, p=0.2063). Groups: Water (n=5 cells from 4 mice; 2 males, 2 females), Nicotine (n=7 cells from 5 mice; 2 males, 3 females). Data shown are from the second minute of baseline. **(C)** Representative traces from adolescent nicotine-treated (red) and adolescent water-treated (black) adult mice before and after bath application of CNO (50 μM). **(D)** There is a main effect of wash-on (F(1, 10) = 39.23, p<0.0001). However, there is no difference in the decrease in spontaneous firing rate between minute 2 in ACSF and minute 11 in CNO among cells from adolescent water-treated and adolescent nicotine-treated mice (two-way RM ANOVA: F(1, 10) = 0.8123, p=0.3886). Sidak’s post hoc analysis revealed that cells from both adolescent water-treated and adolescent nicotine-treated mice decreased their firing between the 2^nd^ minute of baseline in ACSF and minute 11 in CNO (p=0.0112 and p=0.0005, respectively). **(E)** Normalized changes in spontaneous action potential firing between the adolescent nicotine-treated and adolescent water-treated groups in adulthood. Comparing between groups over time, bath application of CNO (50 μM) did not produce a difference in firing rate between groups (two-way RM ANOVA: F(10, 100) = 0.5899, p=0.8188). There is a main effect of Time (F(10, 100) = 15.92, p<0.0001). Data presented as mean ± SEM, *p<0.05, ***p<0.001, ****p<0.0001. P, postnatal day.

**Supplementary Table 1.**
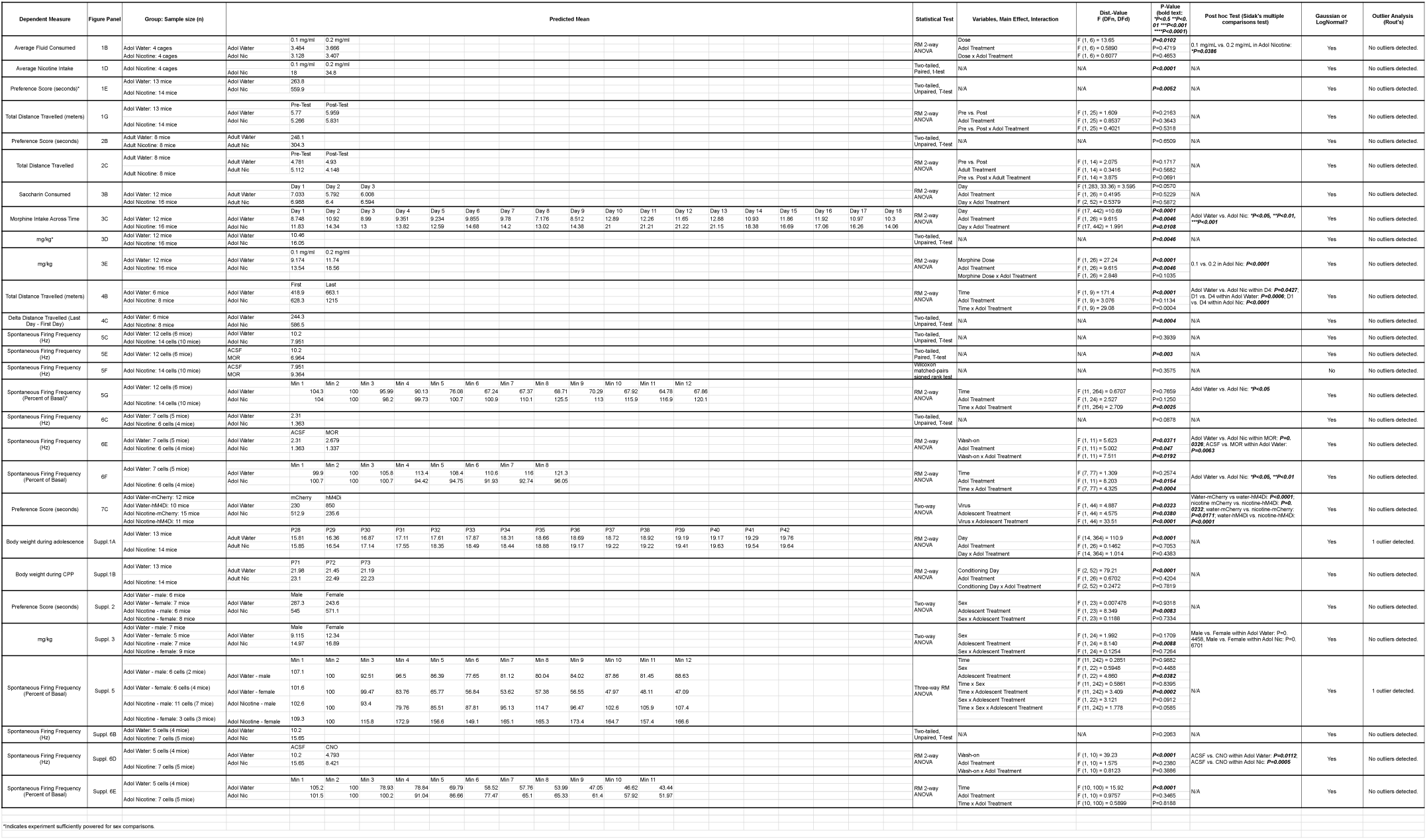
Statistical analyses for each figure panel and all results.

**Supplementary Table 2:**
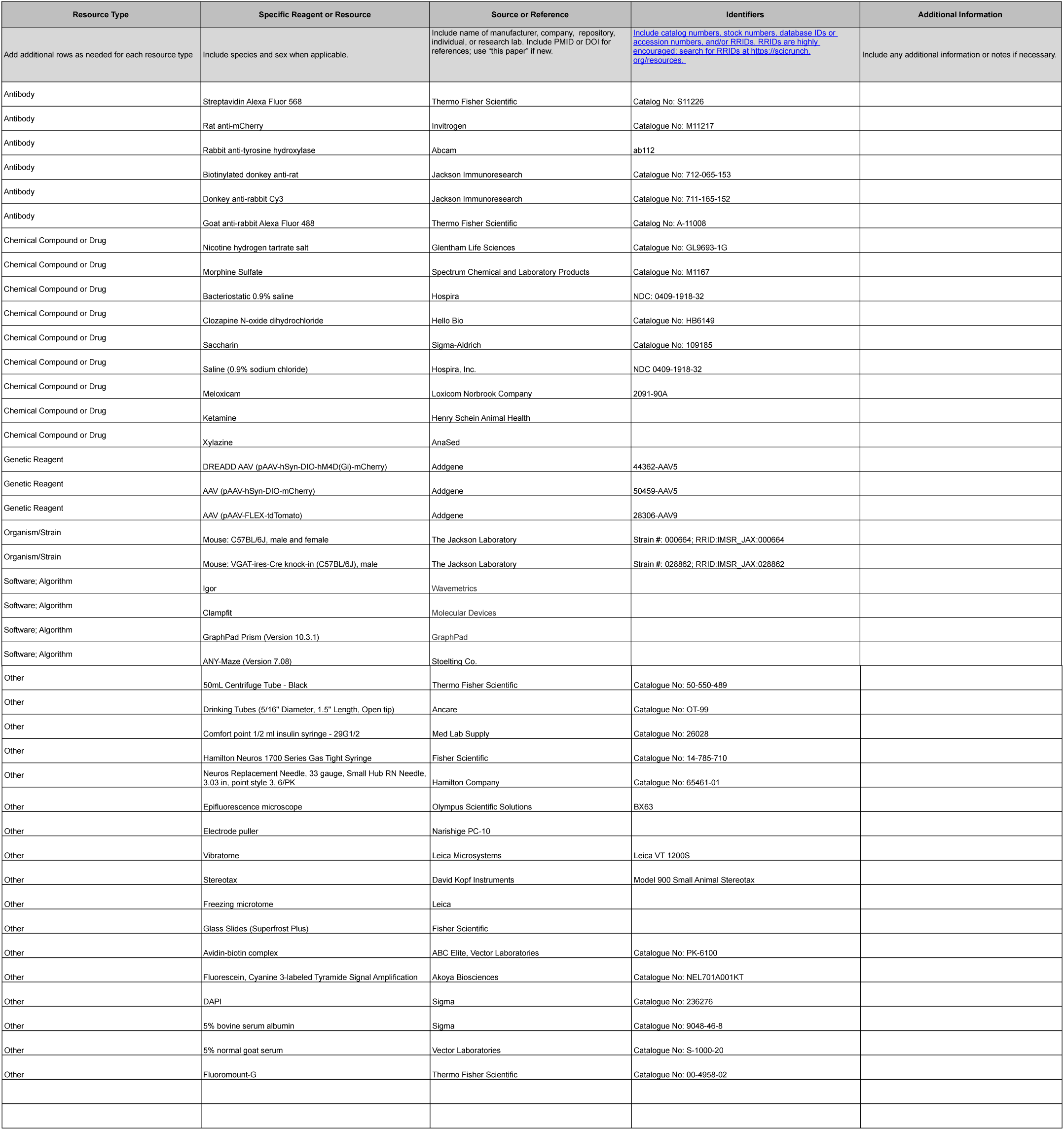
Key Resources Table.

**Supplementary Table 3.**
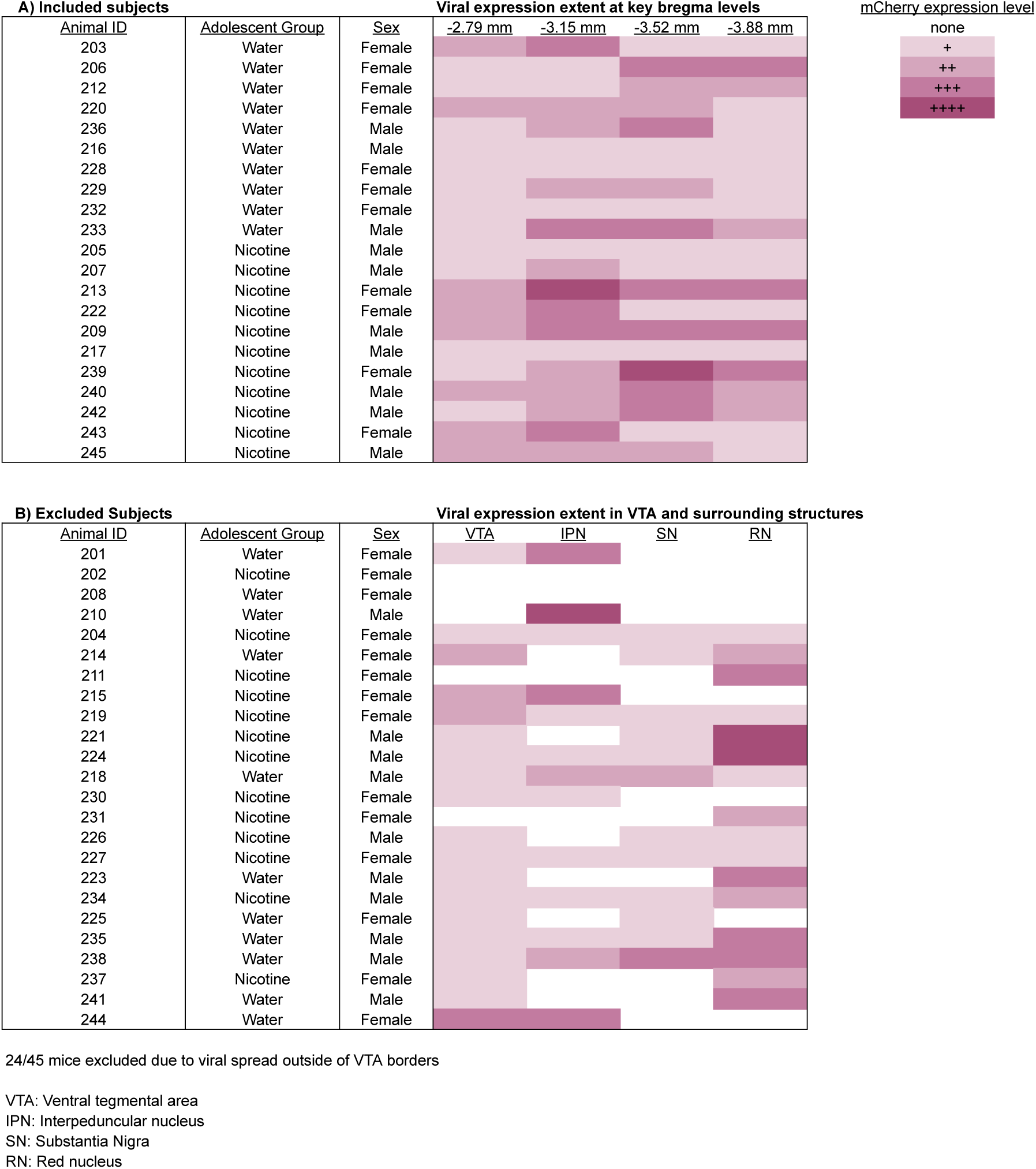
Viral expression in VTA and surrounding structures.

## Notes

### Competing Interest Statement

The authors have declared no competing interest.

